# TGF-beta dynamically controls epithelial identity in a 3D model of human epiblast

**DOI:** 10.1101/2023.12.07.570575

**Authors:** Irene Zorzan, Elena Carbognin, Andrea Lauria, Valentina Proserpio, Davide Benegnù, Caterina Dalrio, Mattia Arboit, Irene Paolucci, Andrea Drusin, Monika Sledziowska, Gianluca Amadei, Salvatore Oliviero, Graziano Martello

## Abstract

Pluripotency is the ability to give rise to all cell types of the body and is first observed in a mass of disorganised cells of the embryo. Upon implantation, pluripotent cells form a columnar epithelium and undergo lumenogenesis. At gastrulation, a portion of the pluripotent epiblast will undergo epithelial to mesenchymal transition (EMT), forming the primitive streak (PS).

It still remains unclear what molecular mechanism supports the epithelial identity of the pluripotent epiblast before gastrulation. Here we developed an optimised, chemically defined 3D model of human pluripotent epiblast formation in which conventional pluripotent stem cells (PSCs) self-organise into a columnar epithelium with a lumen in 48 hours. From 72 hours we observed spontaneous symmetry breaking and specification of PS-like cells, as confirmed by single-cell RNA sequencing.

We found that Insulin and FGF signalling are both required for the proliferation and survival of the pluripotent epiblast model. Conversely, TGF-beta signalling maintains epithelial identity. Epithelial identity appears uncoupled from the expression of canonical pluripotency markers OCT4, NANOG and PRDM14, but under the control of ZNF398. Once the pluripotent epithelium is established, TGF-beta inhibition is inconsequential, and stimulation with Activin A leads to highly efficient PS induction. We conclude that TGF-beta dynamically orchestrates epithelial identity of human pluripotent cells.

## Introduction

Pluripotency is the potential to give rise to all cell types of the adult body and, during development, is first observed in the inner cell mass (ICM) of the pre-implantation blastocyst. The ICM segregates into pre-implantation naive epiblast cells, which retain pluripotency, and hypoblast, or primitive endoderm cells, which will give rise to extraembryonic cells of the parietal and visceral endoderm and tissues like the yolk sack^1–4^. A second population of the pre-implantation blastocyst, the trophectoderm, will give rise to cytotrophoblast, which is crucial for embryo implantation. At the time of implantation, epiblast cells polarise into rosettes leading to the formation of a lumen, which will expand into the future amniotic cavity. The epiblast forms a polarised columnar epithelium, with the apical side facing the amniotic cavity. The epithelial identity of the epiblast is then maintained until the gastrulation stage, as the formation of the primitive streak (PS) entails a transition from epithelial to mesenchymal identity and loss of pluripotency^5^. It still remains unclear whether epithelial identity and pluripotency of the post-implantation epiblast are tightly coupled and whether one affects the other. It is also unclear what signals induce and maintain the epithelial identity in the post-implantation epiblast. The Zernicka-Goetz laboratory reported a critical role for the basal membrane, or basement membrane, produced by the primitive and visceral endoderm and required for the polarization and maturation of the mouse epiblast epithelium^6^. Moreover, remodelling of the basement membrane allows, in mouse embryos, for the expansion of the PS^7^. However, the requirement for a basement membrane does not rule out the presence of instructive signals in this context, in particular in the case of the human post-implantation epiblast, which is understudied for both technical and ethical reasons.

Conventional human pluripotent stem cells (PSCs) are either derived from human embryos as Embryonic Stem Cells (ESCs)^8^, or via reprogramming of somatic cells as induced Pluripotent Stem Cells (iPSCs)^9,10^. Conventional human PSCs are cultured as 2D monolayers and are molecularly close to the post-implantation epiblast^11^. They have been extensively used to study the molecular circuitries controlling the maintenance of pluripotency and their differentiation towards germ layers. Transcription factors including POU5F1/OCT4, SOX2, NANOG, PRDM14 and ZNF398, are either required or sufficient to maintain human pluripotency in PSCs^12–14^. These transcription factors cross-regulate each other, forming a network that is stabilised by signalling via the canonical TGF-beta, FGF and Insulin pathways^13,15–21^.

Culture of PSCs in a 2D monolayer however does not capture the spatial constraints and cell-cell dynamics of the early embryo. To overcome such limitations several 3D models of the human post-implantation embryo have been reported ^22–26,26–32^. These are useful tools for the study of cell-cell communication and self-organisation, which advanced our understanding of some developmental processes. For example, it has been shown that lumenogenesis of the epiblast is controlled by actin cytoskeleton^33^, that amnion specification occurs in response to BMP^22,23,31,32,34^, and that the production of the basement membrane of hypoblast cells is induced by TGF-beta ligands^27^. Across these models, there is a clear trade-off between the complexity (i.e. number of cell types generated) and the efficiency/reproducibility (e.g. fraction of structures correctly organised). Moreover, the presence of multiple cell types and the use of undefined reagents (i.e. conditioned medium, foetal serum or KO serum replacement) might represent a hurdle to identifying the molecules controlling a process of interest.

For these reasons, we optimised a minimal model of the post-implantation epiblast. Under chemically defined conditions, conventional PSCs undergo lumenogenesis and form a polarised columnar epithelium. Epithelial identity is then maintained until a symmetry-breaking event occurs, in which mesenchymal TBXT+ cells are formed. Single-cell RNA sequencing (scRNAseq) confirmed the acquisition of post-implantation epiblast and PS identities.

TGF-beta signalling is crucial during the initial 48 hours for the establishment of epithelial identity, which appears uncoupled from the expression of pluripotency OCT4, NANOG and PRDM14, but under the control of ZNF398. Once the pluripotent epithelium is established, TGF-beta inhibition is inconsequential, while stimulation with the TGF-beta ligand Activin A alone leads to efficient mesoderm induction. We conclude that TGF-beta dynamically orchestrates the epithelial identity of post-implantation pluripotent cells.

## Results

### Efficient generation of a 3D *in vitro* minimal model of epiblast-like structures resembling post-implantation epiblast and primitive streak

After implantation, epiblast cells self-organise in a columnar epithelium that undergoes lumenogenesis, while maintaining pluripotency. It is possible to recapitulate these processes *in vitro* by surrounding hPSCs in an extracellular matrix mimicking the basement membrane^29,32,34–37^. With the aim of studying the mechanisms controlling the epithelial identity of the epiblast, we optimised a chemically defined 3D epiblast model. We first analysed scRNAseq data of human embryos asking what signals the post-implantation epiblast might be responsive to (Extended Data Fig. 1a-b). All 4 FGF receptors (FGFR1/2/3/4), Insulin and IGF1/2 receptors, TGF-beta (TGFBR1/2/3) and Nodal/Activin receptors (ACVR2A/B) were highly expressed in the post-implantation epiblast. Moreover, the ligands FGF2, FGF4, NODAL and TGF-beta 1 were expressed by the post-implantation epiblast; NODAL and IGF1 were highly expressed in the hypoblast, while TGF-beta 1, Activin A and IGF2 were highly expressed by cytotrophoblast cells (Extended Data Fig. 1a-b). We conclude that in the post-implantation epiblast, the TGF-beta, FGF and Insulin/IGF pathways could be active, thus we decided to use a chemically defined medium (E8), which contains ligands of these 3 pathways.

We plated a suspension of single hPSCs on top of a layer of growth factor-reduced Matrigel (GFRM) in E8 medium containing 5% GFRM (Extended Data Fig. 1c). By optimising the number of cells plated, we observed the efficient formation of cell aggregates that after 2 days started to form a lumen and at day 4 appeared as large cysts composed of a columnar epithelium (Fig. 1a). From 50,000 single cells seeded we obtained 11652 +/-1400 cysts at day 4, and >95% displayed the correct formation of a lumen and epithelial organisation (Fig. 1b).

**Figure 1:**
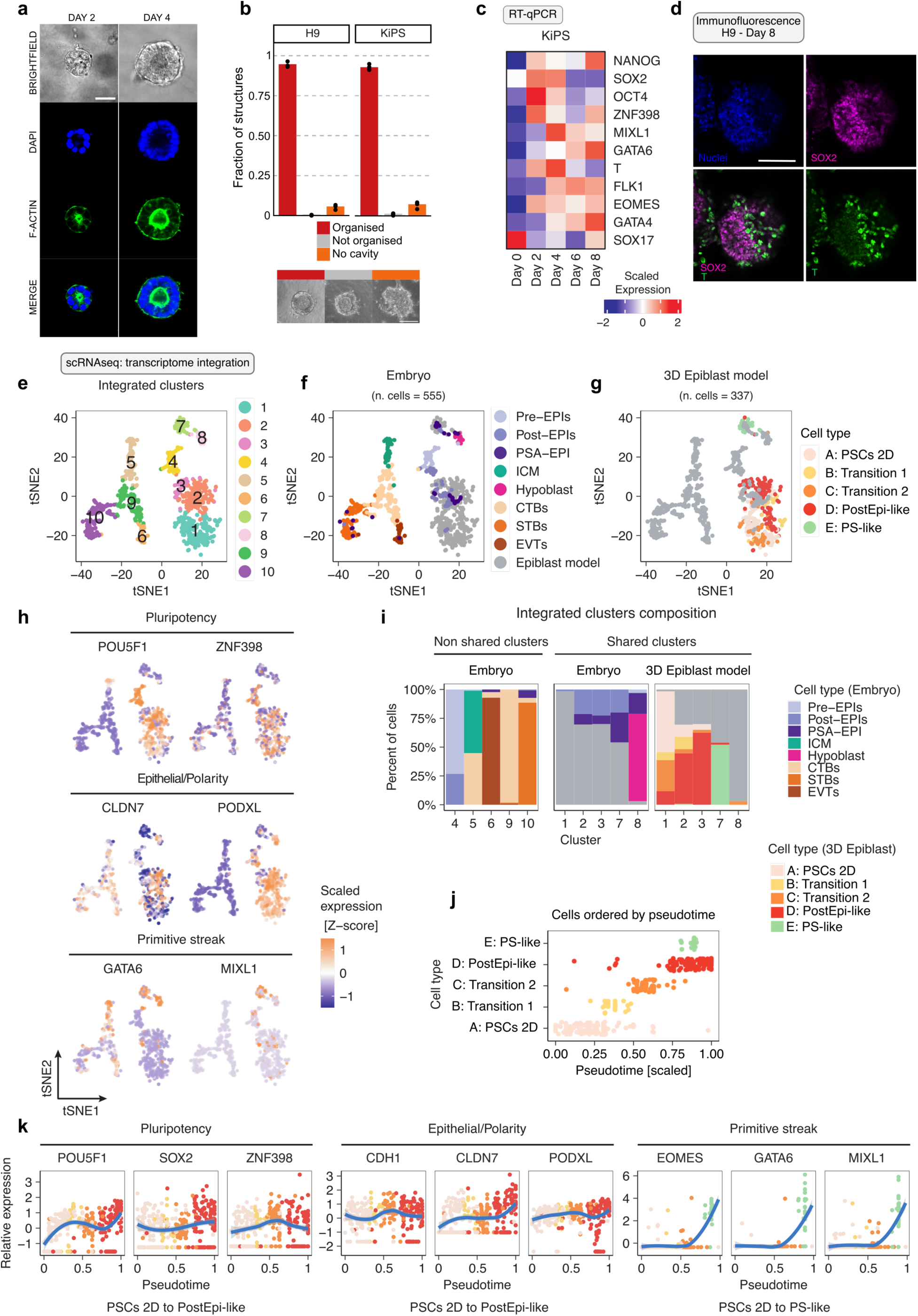
Optimisation and characterisation of a 3D epiblast model. **a,** Left: Representative images and immunostaining for DAPI and Phalloidin of 3D structures after 2 or 4 days of self-organisation. Scale bars 30µm. **b,** Bar plot showing the number of structures obtained after 4 days of self-organisation in two different cell lines, H9 and KiPS respectively. Red bars indicate epithelial organised structures, grey bars indicate not organised structures and orange bars indicate structures without a cavity. Bars indicate the mean of 4 independent experiments, shown as dots. **c,** Heatmap showing gene expression analysis by RT-qPCR for markers of pluripotency and primitive streak in a time-course of 8 days of self-organisation of KiPS cells. Data are presented as scaled log-expression values averaged across 4 independent experiments). Red and blue indicate high and low expression, respectively. **d,** Representative images of immunostaining for SOX2 and T (TBXT, also known as Brachyury) of KiPS (wild-type) after 8 days of self-organisation. On day 8, the 3D structures still express pluripotency markers, such as SOX2, but they also expressed the anterior primitive streak marker, T. Note that SOX2 and T markers are mutually expressed (see merge SOX2/T). Scale bars 50µm. **e, f, g,** t-SNE embedding of 892 single cell transcriptomes showing the results of the comparative integrated analysis of the embryo^41^ and 3D epiblast datasets. Cells are coloured by clusters (E), embryo cell populations (F, number of cells = 555) and 3D epiblast model populations (G, number of cells = 337). **h,** t-SNE embedding of 892 single cell transcriptomes coloured according to the relative (Z-score) expression levels of selected marker genes for pluripotency (POU5F1, ZNF398), polarity (CLDN7, PODXL) and primitive streak (GATA6, MIXL1) cell identities. **i,** Bar plots showing the relative proportions of cell populations in each integrated cluster for the non-shared (left) and shared (right) clusters between the embryo and the 3D epiblast dataset, calculated with respect to their differentiation day and cell type. **j,** Scatter plot showing the pseudo-temporal ordering of cell types of the 3D epiblast model according to their developmental trajectory inferred by Monocle3. X-axis reports pseudotime values rescaled in the 0-1 range, Y-axis reports cell types. **k,** Scatter and line plots showing the gene expression trends across pseudotime of selected marker genes for pluripotency (POU5F1, SOX2, ZNF398) and epithelial/polarity (CDH1, CLDN7, PODXL) identities in the PSCs to PostEpi-like transition (left and middle panels), and primitive streak (EOMES, GATA6, MIXL1) identity in the PSCs to PS-like transition (right panel). X-axis reports pseudotime values rescaled in the 0-1 range, Y-axis reports expression levels, dots are coloured by cell type.

The cysts displayed apical accumulation of F-actin and basal nuclei, as reported ^29,32–34,36^. Up to day 4 the cysts maintained expression of pluripotency markers NANOG, OCT4 and ZNF398 at levels higher than those of conventional hPSCs in 2D (Fig. 1c, Extended Data Fig. 1d).

Pluripotent cells of the epiblast form the PS, a migratory cell population of the human embryo, transiently expressing TBXT (also known as T or BRACHYURY), GATA6, MIXL1 and EOMES^22,40,41^ ultimately giving rise to the mesoderm and endoderm germ layers. We observed robust expression of several PS/mesoderm markers from day 4 onwards (Fig. 1c and Extended Data Fig. 1e), with TBXT+ cells forming clusters (Fig. 1d). These results were unexpected, given that we did not provide exogenous BMP or WNT ligands, which induce PS in several human *in vitro* models^22,23,27,29,32,37^. RNAseq analysis revealed an elevated expression of the ligands WNTA3/5A and BMP2/4/7, and of their respective target genes AXIN1 and ID1/2/3 in day 4 cysts (Extended Data Fig. 1f). These results indicate that autocrine WNT and BMP signalling might be responsible for the induction of PS genes.

Next, we performed scRNAseq^42^ to determine the identity of individual cells forming cystic structures. We analysed the PSCs cultured in 2D used to form 3D structures (day 0) and structures collected at day 2, 4 and 6 (Extended Data Fig. 2a, left).

We identified 6 clusters (Extended Data Fig. 2a, right). Clusters A, B, C and D expressed high levels of pluripotency and epithelial/polarity markers, while cluster E expressed PS markers. Cluster F comprised only cells from day 0 and day 2 (Extended Data Fig. 2b), which expressed only SOX2, but no other pluripotency or PS markers (Extended Data Fig. 2c). We conclude that these cells underwent rapid spontaneous differentiation, they failed to maintain any defined cell identity, and thus we excluded them from further analyses.

Next, we integrated our scRNAseq data with data from human embryos^41^. We identified 10 clusters (Fig. 1e), most of which corresponded to the different cell types previously annotated by Xiang and colleagues (Fig. 1f). For example, cluster 5 (Fig. 1e) corresponded to ICM cells (Fig. 1f). Interestingly, cluster 7 comprised both cells from the early PS of the embryo and cells from our *in vitro* model (Fig. 1f-g). Cluster 7 displayed elevated expression of PS markers GATA6, MIXL1, EOMES, VIM and CDH2 (also known as NCAD or N-Cadherin)^22,27,40,41^ (Fig. 1h and Extended Data Fig. 2d). We conclude that our cystic structures contain cells transcriptionally compatible with PS identity.

Clusters 2 and 3 contained mostly cells of the post-implantation epiblast (Post.EPI) and cells from cysts (Fig. 1i). Cells in clusters 2 and 3 did not express markers of ICM (ESRRB^41^) and Preimplantation Epiblast (TFCP2L1 and KLF17^41,43,44^), nor PS markers, while they did express high levels of general pluripotency (POU5F1, SOX2, PRDM14 and ZNF398) and polarity/epithelial markers (CLDN6/7 and PODXL) which have been detected in the post-implantation epiblast ^27,32,36^ (Fig. 1h and Extended Data Fig. 2e-f). These analyses indicate that our *in vitro* model contains cells transcriptionally compatible with Post-implantation epiblast identity. Finally, cluster 1 of this integrated analysis was entirely composed of cells from our *in vitro* model (Fig. 1i).

Thanks to the integrated analysis we were able to classify the different populations of our 3D model. Cluster A of our *in vitro* model was composed mainly of PSCs at day 0 (Extended Data Fig. 2b), thus named ‘PSC_2D’. Cluster E was the main contributor to cluster 7 of the integrated analyses (Fig. 1i), therefore it was named ‘PS-Like’. Indeed, cluster E displayed a loss of CDH1 expression, a hallmark of PS formation^5,25^. Cluster D contained cells mostly from day 6 (Extended Data Fig. 2b) and was the major contributor to clusters 2 and 3 of the integrated analyses (Fig. 1i, red), therefore it was named ‘PostEpi-Like’. Two clusters (B and C) were mostly composed of cells at day 2, transitioning from PSC in 2D to the PS-like and PostEpi-like states (‘Transition1’ in yellow, ‘Transition2’ in orange).

We then performed pseudotime analysis of the *in vitro* differentiation trajectories and found that pluripotency and epithelial/polarity markers increased in expression going from PSC_2D to Transient1/2, peaking in the PostEpi-like cluster (Fig. 1j-k and Extended Data Fig. 2g). Transition from PSC_2D to PS-like was characterised by a sharp increase in PS markers.

We concluded that our chemically-defined protocol efficiently gives rise to 3D structures that undergo luminogenesis, form a columnar epithelium with Post-implantation epiblast transcriptional identity, eventually specifying PS-like cells. Based on naming conventions recently proposed ^45^, we named our model 3D-hE-gastruloids.

### TGF-beta is required for epithelialisation of 3D-hE-gastruloids

Our 3D minimal model recapitulates lumenogenesis, formation of a pluripotent epithelium and specification of PS-like cells in the absence of extraembryonic cells, under chemically-defined conditions, thus representing an ideal setting to identify the signalling pathways controlling these processes.

We functionally tested the three main signalling pathways (Insulin, FGF and TGF-beta) activated by our chemically-defined medium (Extended Data Fig. 1a-c). First, we plated hPSCs in absence of Insulin and observed widespread cell death, indicating that Insulin is required for cell survival (Fig. 2a).

**Figure 2:**
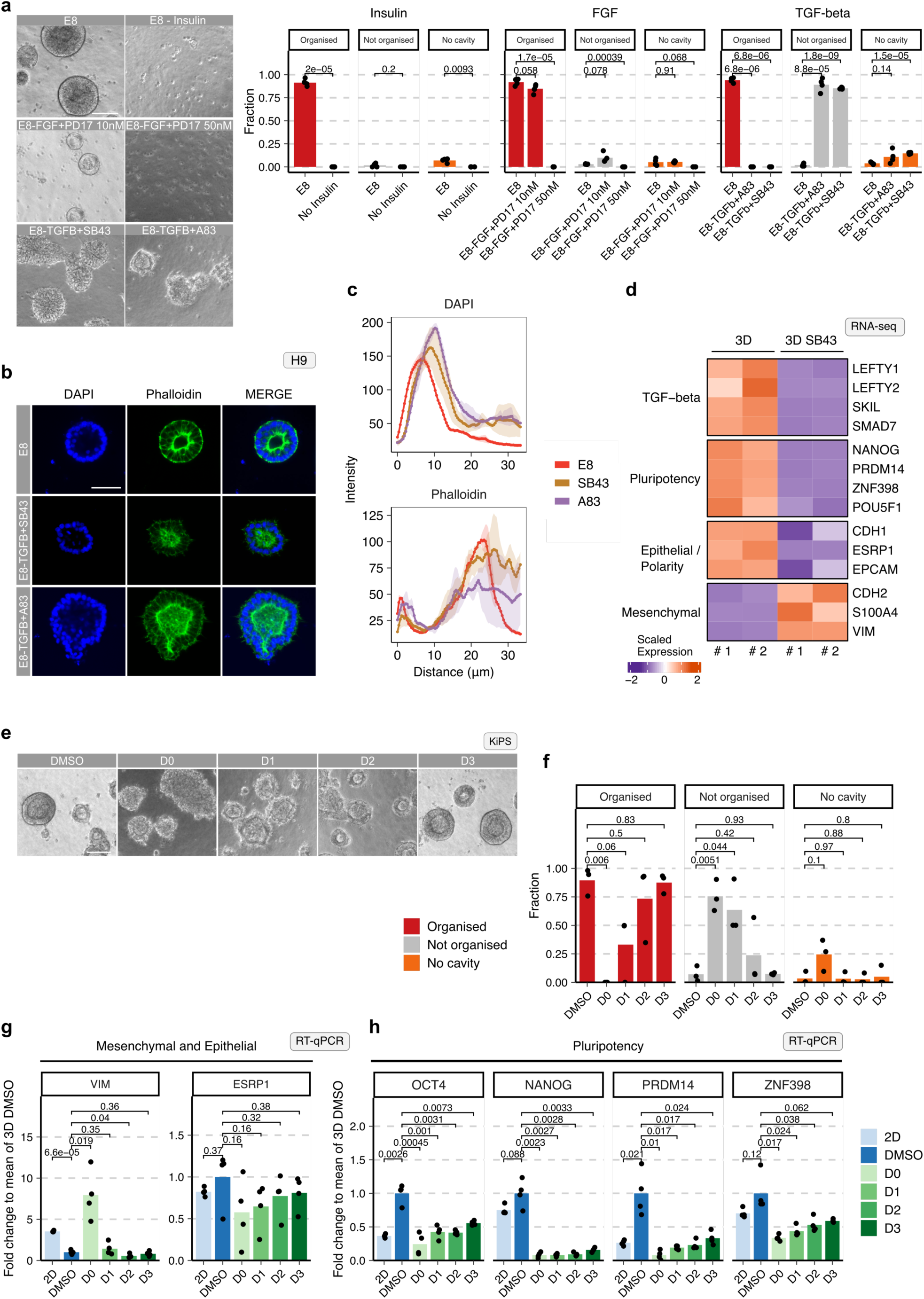
TGF-beta is required for epithelialisation of 3D-hE-gastruloids. **a,** Left: representative images of 3D structures obtained under the indicated conditions. Right: fraction of structures displaying a lumen and a correctly organised epiblast epithelium (Organised - red bars), a lumen with alterations in the epithelium (not organised - grey bars) or absence of lumen (no cavity - orange bars). Bars indicate the mean of 4 independent experiments, shown as dots. Unpaired two-tailed Welch’s *t*-test. **b,** Representative images of immunostaining for Phalloidin and DAPI of 3D structures obtained under the indicated conditions. Scale bar: 50µm **c,** Quantification of Phalloidin and DAPI intensity along a radius connecting the centre of a structure to its edge (see Method details). N=3 structures, for which 7 randomly selected radii were quantified. Line indicates the mean, shaded areas the s.e.m. **d,** Heatmap showing gene expression levels measured by RNA-seq for TGF-beta targets, pluripotency and primitive streak markers that are differentially expressed in 3D structures following TGF-beta inhibition with SB43 as compared to control. Scaled log-expression values from 2 independent experiments. Orange and purple indicate high and low expression, respectively. **e,** Representative images of KiPS-derived structures following a 4 days time-course of TGF-beta inhibition with A83. **f,** Bar plots showing fraction of organised (red bars), not organised (grey bars) and no cavity (orange bars) KiPS-derived structures in a 4 days time-course of TGF-beta inhibition with A83. Unpaired two-tailed Welch’s *t*-test. **g,** Bar plots showing gene expression analysis by RT-qPCR of mesenchymal/epithelial and polarity (left) and pluripotency (right) marker genes performed in KiPS-derived structures following a 4 days time-course of TGF-beta inhibition with A83. Unpaired two-tailed Welch’s *t*-test.

FGF ligands are produced by the epiblast itself (Extended Data Fig. 1b) and also in our 3D structures, thus we tested FGF function by withdrawing from the medium and adding the FGF Receptor inhibitor PD173074 (PD17). We observed reduced size in otherwise correctly formed structures at low doses (Fig. 2a), and decreased survival at higher doses, indicating that FGF induces proliferation and survival, as recently reported for *in vitro* cultured human embryos^43^.

In the case of TGF-beta, withdrawal of the ligand and blockade of TGF-beta signalling with two independent inhibitors, SB431542 (SB43) and A83-01 (A83) had similar effects on self–organisation, without impacting survival (Fig. 2a and Extended Data Fig. 3a-b). While control structures displayed basal nuclei, sharp apical F-actin domain and columnar shape (Fig. 2b-c), TGF-beta inhibition caused a shift in the nuclei position, loss of a defined apical F-actin domain and an irregular shape. Transcriptome analyses revealed a reduction of epithelial markers CDH1, ESRP1 and EPCAM, with an increase in mesenchymal markers CDH2, S100A4 and VIMENTIN (Fig. 2d). Furthermore, a fraction of 3D-hE-gastruloids failed to form a cavity, indicating a partial impairment of lumenogenesis (Fig. 2a and Extended Data Fig. 3a-b). These results indicate that TGF-beta signalling is required to maintain pluripotent cells as a polarised columnar epithelium.

Of note, TGF-beta inhibition led also to downregulation of key pluripotency markers (Fig. 2d), indicating that TGF-beta signalling maintains pluripotency, in agreement with previous studies in 3D and 2D models^13,19,20,31,46^.

Next, we asked when TGF-beta signalling is needed for the correct formation of 3D-hE-gastruloids (Fig. 2e-f and Extended Data Fig. 3c-e). We observed that inhibition of TGF-beta starting at the time of plating single cells (day 0) or at day 1 strongly impaired self-organisation, but was inconsequential if it started at day 2 or 3 (Fig. 2e-f and Extended Data Fig. 3c-e). Consistently, the expression of epithelial and mesenchymal markers was affected only when TGF-beta was inhibited from day 0 or day 1 (Fig. 2g and Extended Data Fig. 3f). Interestingly, pluripotency markers were strongly downregulated even when TGF-beta was inhibited at day 3 (Fig. 2h and Extended Data Fig. 3g). We conclude that maintenance of epithelial identity requires TGF-beta signalling only during the first 2 days, whilst expression of pluripotency markers requires constant TGF-beta stimulation.

### TGF-beta promotes the self-organisation of murine PSCs

Ligands of TGF-beta, like Nodal are highly expressed at the time of implantation in murine embryos, when the epiblast acquires polarity and epithelial identity, forming a rosette and then a lumen. We therefore hypothesised that TGF-beta might also promote the formation and organisation of a pluripotent epiblast epithelium in the mouse. To test this hypothesis we used two *in vitro* model systems.

First, we used mouse naive ESCs, embedded in GFRM in the presence of the basal medium N2B27. These conditions lead to the formation of rosette structures after 96 hours, with basal nuclei and apical accumulation of F-actin^47–49^ (Fig. 3a-b). Inhibition of TGF-beta throughout the process led to the formation of structures in which the epithelial organisation was lost and cavity formation was impaired in ∼30% of the cases (Fig. 3c). Pou5f1, also known as Oct4, is highly expressed in rosettes, like in the peri-implantation embryo^50^, and its expression is reduced by TGF-beta inhibition (Fig. 3d). Second, we combined wild-type mouse ESCs with inducible-Gata4 ESCs in the absence of exogenous extracellular matrix proteins. This system has been shown to self-organise into structures with an inner epiblast-like layer of pluripotent, OCT4 positive cells, surrounded by a layer of visceral endoderm (VE)-like cells^51^ (Fig. 3e-f). Accumulation of F-actin and Podocalyxin strongly marks VE-like cells as well as the lumen-facing, apical side of the epiblast-like cells (Fig. 3f). Inhibition of TGF-beta completely ablated lumen formation and epithelial organisation. F-actin and Podocalyxin staining was retained only by VE-like cells, but was not detected in the core of the structures, indicating impaired polarisation. The expression of Oct4 was dramatically reduced both at the protein and mRNA levels. We conclude that TGF-beta is required for the correct organisation of a pluripotent epithelium and for maintenance of the pluripotency marker Oct4, as also observed in our human PSC-based model.

**Figure 3:**
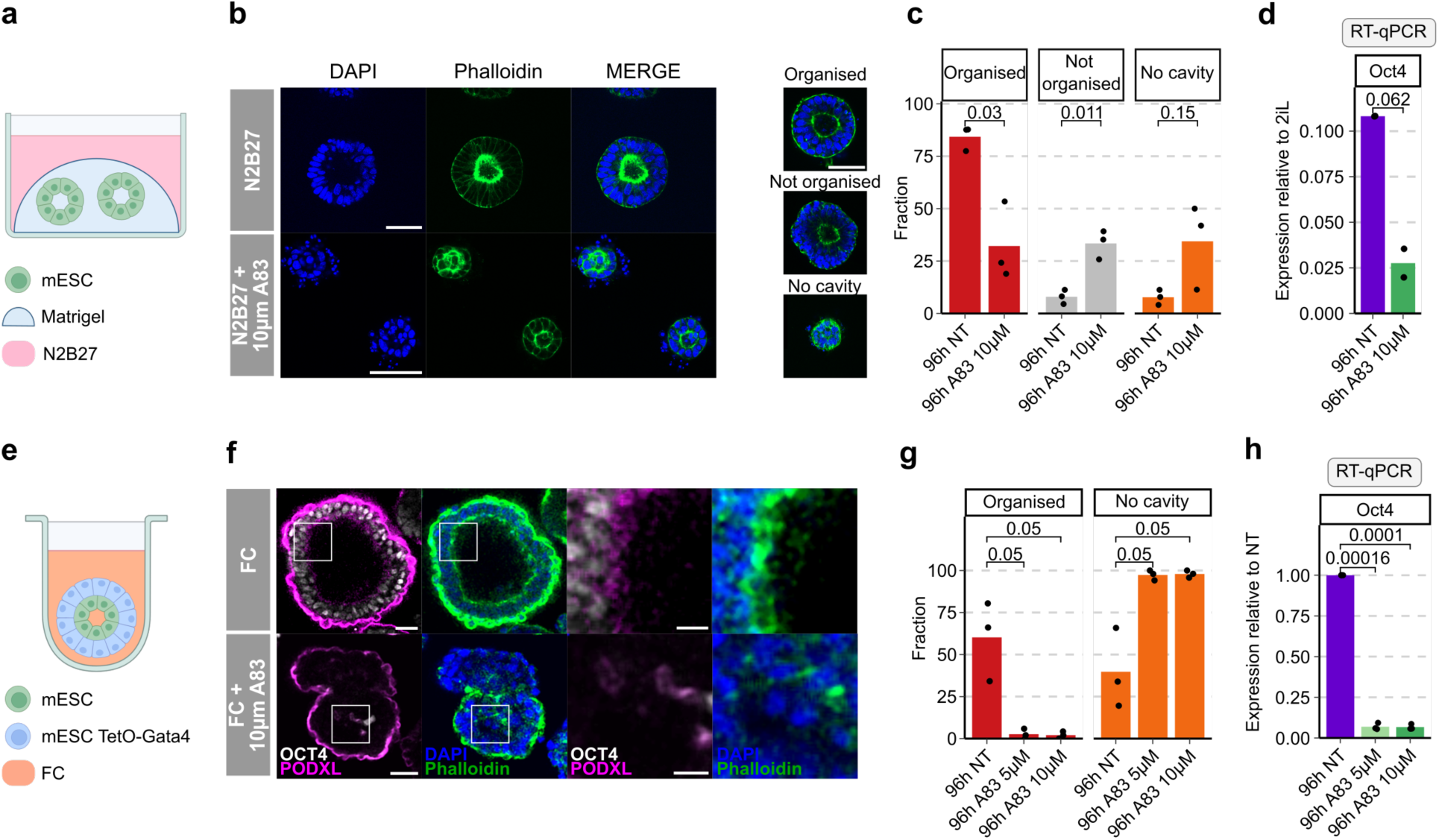
TGF-beta promotes the self-organisation of murine PSCs. **a,** Schematic representation of the experimental assay used to generate rosette-like structures with mESCs. **b,** Representative images of rosette-like structures from mouse naive ESCs, cultured for 96 hours in N2B27 or in the presence of A83 inhibitor. Nuclei were stained with DAPI, f-actin was stained with Phalloidin. Right: examples of morphology of rosette-like structures treated with A83 inhibitor that display either an epithelial organisation (organised), a disorganisation (not organised), or the absence of a cavity (no cavity). Scale bar: 50µm. **c,** Bar plot showing the number of rosette-like structures obtained after 96 hours, with or without (NT) treatment with A83. Red bars indicate epithelial organised structures, grey bars indicate not organised structures, and orange bars indicate structures without a cavity. Bars indicate the mean of 3 independent experiments, shown as dots. **d,** Bar plots showing gene expression analysis by RT-qPCR of Oct4 in rosette-like structures after 96h of treatment with A83. Unpaired two-tailed Welch’s t-test. **e,** Schematic representation of the experimental assay used to generate cysts with a combination of wild-type mESCs and inducible-Gata4 mESCs. **f,** Representative images of cysts, generated with a combination of wild-type mESCs and inducible-Gata4 mESCs, cultured for 96 hours in Fc medium or in the presence of A83 inhibitor. Nuclei were stained with DAPI, f-actin was stained with Phalloidin. Scale bar: 30µm. Scale bar zoom: 10µm. **g,** Bar plot showing the number of cysts obtained after 96 hours, with or without (NT) treatment with A83. Red bars indicate epithelial organised structures, grey bars indicate not organised structures, and orange bars indicate structures without a cavity. Bars indicate the mean of 3 independent experiments, shown as dots. **h,** Bar plots showing gene expression analysis by RT-qPCR of Oct4 in cysts after 96h of treatment with two doses of A83. Unpaired two-tailed Welch’s t-test.

### ZNF398 promotes epithelial identity downstream of TGF-beta

We previously reported that ZNF398 induces pluripotency and epithelial identity downstream of the TGF-beta pathway in hPSCs in 2D^13^, thus we wondered whether it could also play a role during the self-organisation of hPSCs in 3D. We first measured the levels of ZNF398 in 3D and observed a strong reduction in the presence of SB43, indicating that TGF-beta maintains ZNF398 expression (Fig. 4a and Extended Data Fig. 4a, blue and yellow bars). SB43 also blocked the organisation of 3D-hE-gastruloids (Fig. 2 and 4b-c), causing a reduction of epithelial markers and an increase in mesenchymal ones (Fig. 4d and Extended Data Fig. 4a). We checked the expression of the TGF-beta targets (LEFTY1, LEFTY2) and confirmed robust inhibition of TGF-beta signalling by SB43 in both cell lines (Fig. 4e). We conclude that reduced levels of ZNF398 are correlated with loss of epithelial identity in 3D.

**Figure 4:**
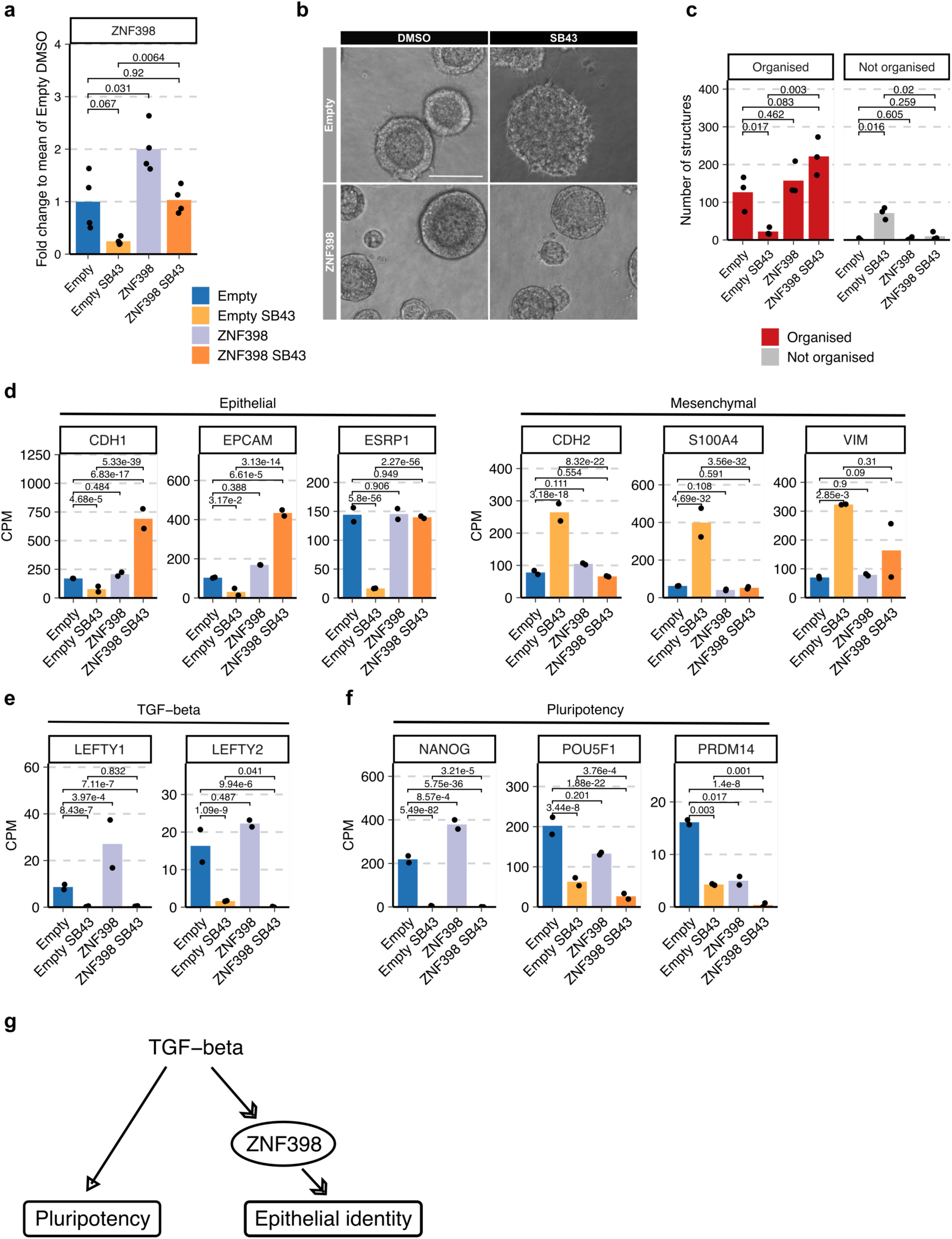
ZNF398 promotes epithelial identity in a 3D epiblast model. **a,** Bar plot showing the expression of ZNF398, measured by qPCR in KiPS stably expressing the empty vector control (Empty) without (blue bar) or with SB43 (yellow bar), or expressing ZNF398 without SB43 (lilac bar) or with SB43 (orange bar) after 4 days of self-organisation. Bars indicate the mean of 4 independent experiments, shown as dots. Unpaired two-tailed Welch’s *t*-test. **b,** Representative morphology of KiPS stably expressing the empty vector (Empty) or ZNF398 in presence of DMSO or SB43. SB43 abolishes the formation of epithelial organisation, which are rescued by forced expression of ZNF398. Scale bars 100 µm. **c,** Bar plot showing the number of structures obtained after 4 days of self-organisation. Red bars indicate epithelial organised structures, while grey bars indicate disorganised structures. Bars indicate the mean of 3 independent experiments, shown as dots. *P*-values calculated by Quasi-Poisson regression of count data. **d, e, f,** RNAseq of KiPS stably expressing the empty vector control (Empty) without (blue) or with SB43 (yellow), or expressing ZNF398 without SB43 (lilac) or with SB43 (orange) after 4 days of self-organisation. Bars indicate the mean of 2 independent experiments, shown as dots. *P*-values calculated by *DESeq2* (see Methods). **g,** Schematic representation of the proposed regulatory module in which ZNF398 promotes epithelial identity downstream of TGF-beta signalling.

To test whether ZNF398 is actively controlling epithelial identity, we generated hPSCs expressing ZNF398 at ∼endogenous levels (Fig. 4a and Extended Data Fig. 4a, lilac bars). We observed formation of correctly epithelialized 3D-hE-gastruloids in the ZNF398 expressing cell line both in the absence and in the presence of SB43 (Fig. 4b-c). Epithelial markers were highly induced while mesenchymal markers were kept at low levels by ZNF398 forced expression (Fig. 4d and Extended Data Fig. 4a), indicating that ZNF398 is epistatic to TGF-beta signalling.

Analysis of a large set of human pluripotency markers (Fig. 4f and Extended Data Fig. 4b) revealed that ZNF398 did not maintain their expression in presence of SB43, indicating that induction of epithelial genes by ZNF398 does not require the presence of other pluripotency factors. We conclude that ZNF398 is sufficient to maintain the epithelial identity in 3D-hE-gastruloids, acting downstream of TGF-beta (Fig. 4g).

### Activin boosts the primitive streak differentiation in the 3D structures

Our results indicate that TGF-beta signalling is required during the first 2 days of formation of 3D structures to acquire a stable epithelial identity (Fig. 2e-g and Extended Data Fig. 3d-g). However, TGF-beta signalling is also known to induce PS and mesendoderm specification in several model systems, including human PSCs. Given that PS formation requires acquisition of a mesenchymal phenotype, it might appear as if TGF-beta promotes, counterintuitively, both epithelial and mesenchymal identities.

We reasoned that these two processes are temporally separated during development, as the epiblast forms a columnar epithelium at the time of implantation, while a subset of epiblast cells undergo EMT and form a PS later, at gastrulation. Given that 3D-hE-gastruloids recapitulate both processes, we decided to exploit it to study TGF-beta responsiveness at different times. As PS-like cells of 3D-hE-gastruloids expressed MIXL1 (Fig. 1k and Extended Data Fig. 2c), we employed a MIXL1-GFP HES3 reporter hESC line.

We allowed for the acquisition of epithelial identity in E8 for 2 days (Fig. 5a). On day 2, structures were either kept in E8 or exposed to the basal medium E6, which lacks FGF2 and TGFB1. Both conditions led to a similar induction of MIXL1, GATA6 and TBXT mRNAs and of the GFP reporter (Fig. 5b-c). We concluded that FGF2 and TGFB1 do not play a role in PS-like cell induction.

**Figure 5:**
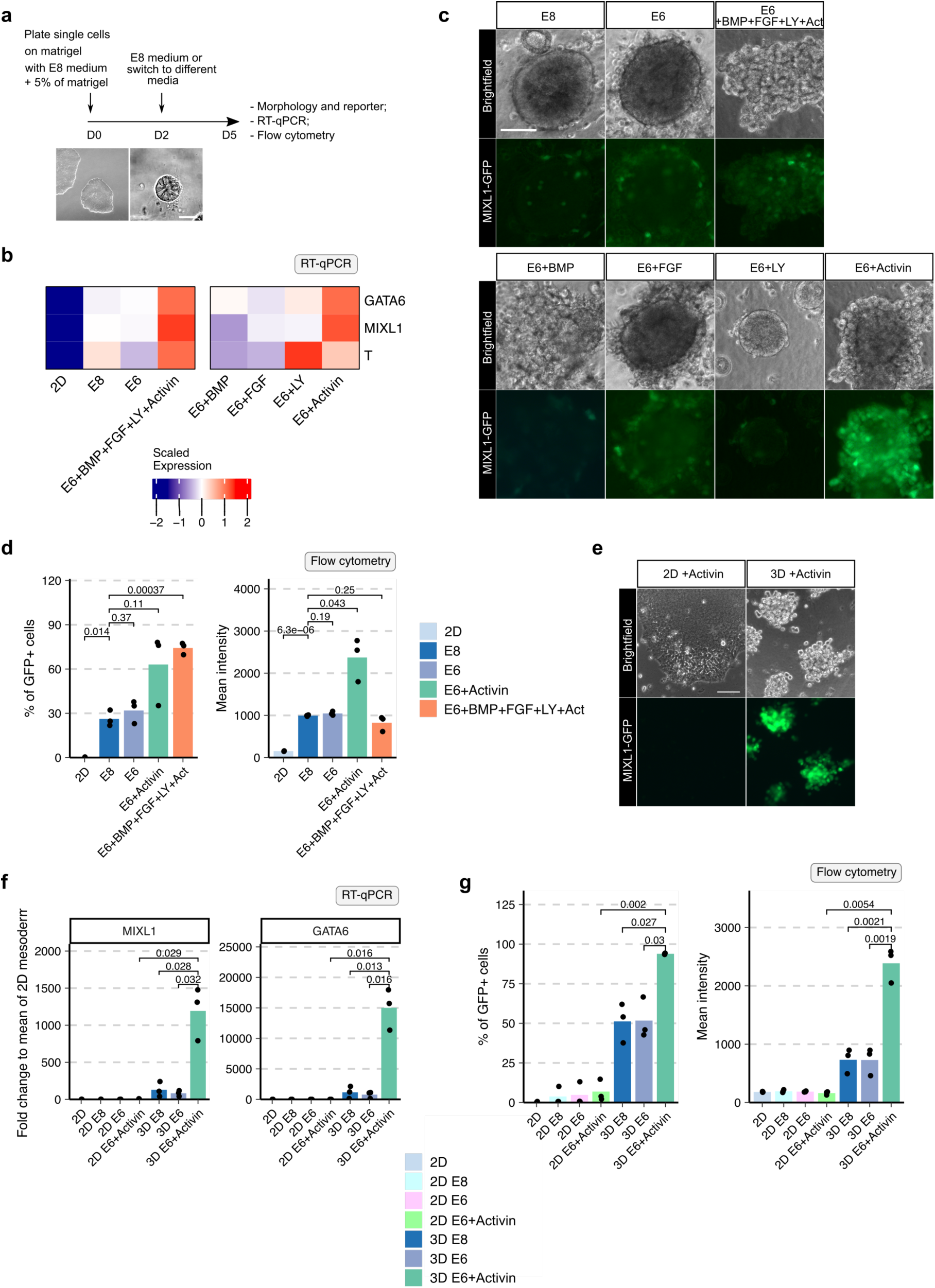
Activin boosts the primitive streak differentiation in the 3D structures. **a,** Schematic representation of the experimental strategy: single hPSCs were seeded on a Matrigel layer with E8 medium + 5% of Matrigel (v/v) at day 0. After 2 days, media was changed with E8 or other different media (E6, E6+BMP+FGF+LY+Activin, E6+BMP, E6+FGF, E6+LY, E6+Activin). On day 5, 3D structures were analysed as indicated. Representative images at day 0 and at day 2 are shown. Scale bar: 30µm. **b,** Heatmap showing gene expression analysis by qPCR for markers of primitive streak in the indicated conditions. Values are expressed as fold-change relative to the mean of E8 samples. Red and blue indicate high and low expression, respectively. **c,** Representative bright field images and fluorescence images of MIXL1-GFP reporter for the indicated conditions at day 5 are shown. Representative images of 4 independent experiments are shown. Scale bar: 50µm **d,** Percentage of GFP positive cells (left panel) and mean intensity of GFP (right panel) analysis by flow cytometry in the indicated conditions. Bars indicate the mean of 3 independent experiments, shown as dots. Unpaired two-tailed Welch’s *t*-test. **e,** Representative images and fluorescence images of MIXL1-GFP reporter for the indicated conditions are shown. Representative images of 3 independent experiments are shown. Scale bar: 50µm **f,** Bar plots showing gene expression analysis by RT-qPCR of primitive streak marker genes in the conditions at day 5. Unpaired two-tailed Welch’s *t*-test. **g,** Percentage of GFP positive cells (left panel) and mean intensity of GFP (right panel) analysis by flow cytometry in the indicated conditions. Bars indicate the mean of 3 independent experiments, shown as dots. Unpaired two-tailed Welch’s *t*-test.

Activin A is a TGF-beta ligand commonly used to induce PS and mesendoderm from hPSCs, in combination with BMP and FGF ligands and LY294002 (LY), an inhibitor of Phosphatidylinositol 3-Kinase^18,52–54^. Exposure to the 4 signalling molecules led to upregulation of PS markers and complete switch to a mesenchymal morphology (Fig. 5b-c). We then tested the contribution of each signalling molecule and surprisingly the sole stimulation with Activin A was sufficient to strongly induce PS-like cells, while the BMP, FGF and LY had no effect. We quantified the activity of MIXL1-GFP reporter by flow cytometry and observed strong induction in >60% of cells in the presence of Activin A (Fig. 5d).

Several previous studies reported that activation of the TGF-beta pathway is not sufficient to induce PS/mesoderm in human PSCs in 2D^18,53,55,56^, in stark contrast with our 3D results. We performed side-by-side experiments in which MIXL1-GFP hESCs were plated on the same substrate (GFRM) in either 2D or 3D, and then subjected to identical signalling environment (E8 for 2 days followed by E6+Activin A for 3 days). Crucially, we confirmed that PSCs in 2D failed to induce MIXL1 or GATA6, while in 3D we observed some spontaneous activation in E6 that was maximised by Activin A treatment (Fig. 5e-g).

In sum, we conclude that 3D-hE-gastruloids, once allowed to self-organise for 2 days into a polarised epithelium, are highly responsive to Activin A stimulation for induction of PS-like cells.

## Discussion

Pluripotent cells of the epiblast acquire epithelial identity at the time of implantation and maintain it for several days. What factors are required for epithelial identity of pluripotent cells? Are pluripotency and epithelial identity of the epiblast coupled, or can one exist without the other?

A basement membrane is structurally required by all epithelia, indeed *in vitro* models of 3D epiblasts contain either substrates mimicking a basement membrane or hypoblast cells that surround epiblast cells and produce components of the basement membrane ^6,22,23,26,27,29,31–36,57^. However, our results indicate that additional instructive cues are needed, as hPSCs cannot form a correctly organised epiblast epithelium in the absence of TGF-beta signalling, even though a basement membrane is provided. Thus TGF-beta actively stabilises epithelial identity, as we previously reported for hPSCs in 2D^13^.

Interestingly, 2 days of TGF-beta signal are sufficient to stabilise the epithelial identity. In contrast, the expression of key pluripotency factors OCT4, SOX2, NANOG and PRDM14 requires constant TGF-beta stimulation. If TGF-beta signal is present only for the first 2 days, we obtained structures with full epithelial character that lost expression of pluripotency markers. In other words, the epithelial identity is maintained for several days even in the absence of key pluripotency factors.

We also observed that ZNF398 is sufficient to induce epithelial identity downstream of TGF-beta. Indeed, hPSCs stably expressing a ZNF398 transgene can form, in the absence of TGF-beta, 3D structures with stable epithelial identity, while key pluripotency factors are not expressed. Based on these observations we conclude that epithelial identity and pluripotency are two separate, uncoupled processes, as cells can lose pluripotency while maintaining their epithelial character.

In the gastrulating embryo, some epiblast cells form the PS and lose both epithelial identity and pluripotency. However, epiblast cells not contributing to the PS will default towards ectoderm lineage, losing pluripotency while maintaining epithelial identity, indicating that also during embryonic development the two processes are uncoupled.

Epithelial identity appears to be stable once established, unless an external cue induces an EMT. Conversely, pluripotency needs constant support by external cues. These two behaviours underlie different needs of the developing embryo: pluripotency is an intrinsically unstable state that should be rapidly extinguished upon germ layer specification, while epithelial identity is found in several tissues and should be maintained throughout development and in the adult.

TGF-beta signalling is known to induce EMT^58^ and PS, as we also observed in our model. How can the same signalling pathway stabilise epithelial identity and induce EMT in the same system?

First, we observed a temporal gap of 2 days between the two responses. The fact that the same signal might exert opposite functions has precedents during embryonic development, as WNT signalling switches from a posteriorising to an anteriorising signal during frog development in less than 24 hours^59^.

Second, we observed that TGFB1 ligand promotes epithelial character but does not affect PS and EMT induction, while Activin A strongly promotes PS and loss of epithelial identity. Although TGFB1 and Activin A should activate the same downstream mediators (i.e. SMAD2 and SMAD3), they might do so with different intensities and durations. For instance, the Siggia and Brivanlou laboratories showed in human 2D gastruloids, in which PSCs form spatially defined territories of embryonic and extraembryonic cell populations in response to extracellular cues^60^, that a transient SMAD2 activation is sufficient to induce pluripotency genes, while a more prolonged activation is needed for mesendodermal differentiation^46^.

Third, SMAD2/3 can interact with different transcription factors in a stage-dependent manner, like OCT4 in PSCs and FOXH1 in mesodermal precursors^61^, resulting in the regulation of different transcriptional programs.

We observed a strongly increased responsiveness to Activin A in 3D conditions compared to 2D, in terms of induction of PS/mesoderm. Also in a microfluidics-based 3D epiblast model^32^ Activin A was sufficient to induce PS-like cells, while in several 2D models different TGF-beta ligands failed to do so^18,46,53,55,56^.

2D models do not recapitulate the 3D spatial organisation of the embryo (e.g. formation of lumen) and display some features that may not be observed *in vivo*. For example, receptors of the TGF-beta pathway are basolateral localised in PSCs and in the epiblast. By culturing PSCs in 2D, the edge of a colony is more responsive to TGF-beta ligands^46,60^, while in the embryo the basal side of the whole epiblast epithelium is in contact with a source of TGF-beta, the hypoblast. Therefore we would speculate that in 3D epiblast models, the basolateral receptors are more exposed to external signals than under 2D conditions, potentially explaining the increased responsiveness. Future studies will be needed to test the responsiveness and signalling dynamics in 2D vs 3D conditions.

From a purely technical perspective, 3D-hE-gastruloids offer several advantages, as they model epithelial formation, lumenogenesis and primitive streak induction under chemically defined conditions. The absence of extraembryonic cell populations overcomes potential ethical limitations. The assay does not require bio-engineerging devices, but only commercially available reagents, thus favouring reproducibility. Finally, its high efficiency and throughput makes it ideal for the dissection of molecular mechanisms and for global analyses like metabolomics and proteomics.

In conclusion, 3D-hE-gastruloids recapitulate several aspects of post-implantation epiblast development, despite the absence of extraembryonic cells, indicating that a basement membrane and the triple stimulation of FGF, Insulin and TGF-beta pathways are the minimal requirements to consolidate post-implantation epiblast identity and to kickstart PS induction.

## METHOD DETAILS

### Cell Culture

Human primed hiPSCs (KiPS^13,62^, Keratinocytes induced Pluripotent Stem Cells) and hESCs (H9 and HES3-MIXL1-GFP^63^) were cultured in Feeder-free on pre-coated plates with 0.5% growth factor-reduced Matrigel (CORNING 356231) (vol/vol in PBS with MgCl2/CaCl2, Sigma-Aldrich D8662) in E8 medium (made in-house according to^39^) at 37°C, 5% CO_2_, 5% O_2_. Cells were passaged every 3-4 days at a split ratio of 1:8 following dissociation with 0.5 mM EDTA (Invitrogen AM99260G) in PBS without MgCl2/CaCl2 (Sigma-Aldrich D8662), pH8. The H9 line (WA09) was obtained from and used under authorisation from WiCell Research Institute. The HES3-MIXL1-GFP line was generated by gene targeting ^63^ and kindly provided by Prof. Andrew Elefanty. KiPS line was derived by reprogramming of commercial human keratinocytes (Invitrogen) with Sendai viruses encoding for OSKM ^62^ and kindly provided by Austin Smith’s laboratory. All cell lines were mycoplasma-negative (Mycoalert, Lonza).

### Formation of 3D-hE-gastruloids

hPSCs were detached as single cells using TrypLE^TM^ Select (1X) (Thermo Fisher 12563-029). A well of a Chamber slide was covered (Thermo scientific 177402, 0.8 cm^2^ each) with 40µl of pure ice-cold growth factor reduced Matrigel (CORNING Cat. 356231) and incubated for 2 minutes (min) at 37°C to allow Matrigel to solidify. 50,000 cells (for each well of a chamber slide) were resuspended in 250μl of E8 medium with 10µM ROCKi. 10 min after plating, the medium was removed and replaced with 250µl of E8 + 10µM ROCKi containing 5% Matrigel. After 24 hours, the medium was changed (E8 medium containing 5% of Matrigel v/v). The 3D structures were grown at 37°C, 5% CO_2_, 21% O_2_. For the experiments in which we functionally tested the three main signalling pathways, we treated the 3D structures from day 0 with (E8 or E8 plus DMSO) or without insulin (E8-insulin), or without FGF in the E8 medium, plus PD173074 (10nM or 50nM), or without TGF-beta in the E8 medium plus SB431542 at the indicated concentrations (1 µM, 2.5µM, 5µM or 10 µM), or without TGF-beta in the E8 medium plus A83-01 at the indicated concentrations (1 µM, 2.5µM, 5µM or 10 µM). For the time course experiments, we treated the 3D structures with SB43 or A83 at the indicated time points (day 0, day 1, day 2, day3) for 4 days. For the experiments with MIXL1-GFP HES3 reporter cells, after 2 days in E8 media, the media was changed with E8 or in the indicated conditions (E6, E6+BMP+FGF+LY+Activin, E6+BMP, E6+FGF, E6+LY, E6+Activin).

### Formation of cysts

To prepare the AggreWell plate (STEMCELL Technologies 34415), 500ml of anti-adherence rinsing solution (STEMCELL Technologies 07010) was added to each well. The plate was then centrifuged at 1,000 x g for 5 minutes and incubated at room temperature until cells were ready for seeding. Rinsing solution was then aspirated and each well was washed with 1 mL of PBS. PBS was then aspirated and 500 uL of FC media were added to each well. To prepare ESCs for generating cysts, Doxycycline (1 mg/mL) (Sigma-Aldrich D9891-5G) was added to CAG-GFP tetO-Gata4 ESCs 6 hours prior to plating in AggreWell. Both CAG-GFP tetO-Gata4 and E14 ESCs were washed once with 1x PBS, and dissociated with Accutase for 4 minutes at 37 °C. The reaction was stopped by adding 2 mL of feeder cell medium (FC, see below). Cells were dissociated gently by pipetting for 4-5 times and centrifuged at 200 x g for 4 minutes. The cell pellet was washed once with 1x PBS, centrifuged again and resuspended in 1-2 mL of FC. Cell suspensions containing 12,000 Doxycycline-treated CAG-GFP tetO-Gata4 ESCs and 12,000 E14 ESCs were pelleted again by centrifugation. This is the number of cells needed for 1 experimental well and it is scaled linearly for each additional well. After resuspending in 1ml of FC medium, the cell suspension was added dropwise to the AggreWell and the plate was centrifuged at 100 x g for 3 minutes. Media change was performed every 24 hours by adding 1 mL of fresh FC medium after removing 1ml of medium from each well. TGF-β inhibition was performed by adding at the indicated concentrations (5 µM or 10 µM) for the entire culture duration. Cysts were collected at 96 hours post-seeding, and either processed for RNA extraction or fixed and processed for immunofluorescence. Feeder cell (FC) medium comprises Dulbecco’s modified essential medium (Gibco 41966052), 15% foetal bovine serum (Gibco 10270106), 1mM sodium pyruvate (Gibco 11360039), 2mM GlutaMAX (Gibco 35050038), 1% MEM non-essential amino acids (Gibco 11140035), 0.1mM 2-mercaptoethanol (Gibco 31350010) and 1% penicillin/streptomycin (Gibco 15140122).

### Generation of hPSCs stably expressing ZNF398

Stable transgenic hPSCs expressing ZNF398 were generated as previously described in ^13^. hPSCs were transfected with PiggyBac (PB) transposon plasmid with piggyBac transposase expression vector pBase. In order to generate the PB plasmids, the candidate ZNF398 was amplified from cDNA and cloned into a pENTR2B donor vector. Then, the transgene was Gateway cloned into the same destination vector containing PB-CAG-DEST-bghpA and pGK-Hygro selection cassette.

For DNA transfection, 250,000 hPSCs were dissociated as single cells with TrypLE (Gibco 12563-029) and were co-transfected with PB constructs (550 ng) and pBase plasmid (550 ng) using FuGENE HD Transfection (Promega E2311), following the protocol for reverse transfection. For one well of a 12-well plate, we used 3.9 μl of transfection reagent, 1 μg of plasmid DNA, and 250,000 cells in 1 ml of E8 medium with 10 µM Y27632 (ROCKi, Rho-associated kinase (ROCK) inhibitor, Axon Medchem 1683). The medium was changed after overnight incubation and Hygromycin B (200 μg/ml; Invitrogen 10687010) was added after 48 hours. For the overexpression experiments, hPSCs stably expressing an empty vector or ZNF398 were plated to form the 3D structures and treated with DMSO or 10 µM SB43 for 4 days.

### RNA sequencing and analysis

Quant Seq 3’ mRNAseq Library Prep kit (Lexogen) is used for library construction.

Library generation is initiated by oligodT priming. The primer already contains Illumina-compatible linker sequences. After first strand synthesis the RNA is removed and second strand synthesis is initiated by random priming and a DNA polymerase. The random primer also contains Illumina-compatible linker sequences. Second strand synthesis is followed by a magnetic bead-based purification step. The library is then amplified, introducing the sequences required for cluster generation. External barcodes are introduced during the PCR amplification step. Library quantification is performed by fluorometer (Qubit) and bioanalyzer (Agilent).

QuantSeq Forward contains the Read 1 linker sequence in the second strand synthesis primer, hence NGS reads are generated towards the poly(A) tail and directly correspond to the mRNA sequence. QuantSeq FWD maintains strand-specificity and allows mapping of reads to their corresponding strand on the genome, enabling the discovery and quantification of antisense transcripts and overlapping genes. Sequencing is performed on NextSeq500 ILLUMINA instruments to produce 5 million reads (75bp SE) for the sample.

The reads were trimmed using BBDuk (BBMap v. 37.87), with parameters indicated in the Lexogen data analysis protocol. After trimming, reads were aligned to the Homo sapiens genome (GRCm38.p13) using STAR (v. 2.7.6a). The gene expression levels were quantified using featureCounts (v. 2.0.1). Genes were sorted removing those that had a total number of counts below 10 in at least 3 samples. All RNAseq analyses were carried out in the R environment (v. 4.0.0) with Bioconductor (v. 3.7). We computed differential expression analysis using the DESeq2 R package (v. 1.28.1)^64^. DESeq2 performs the estimation of size factors, the estimation of dispersion for each gene and fits a generalised linear model. Transcripts with absolute value of log2[FC] > 1 and an adjusted p-value < 0.05 (Benjamini–Hochberg adjustment) were considered significant and defined as differentially expressed for the comparison in the analysis.

### Single cell RNA library preparation and sequencing

For single cell library prep, ∼30 3D cysts structures for each time point were collected and dissociated using Trypsin/EDTA for 5 min at room temperature. Cells were then washed with PBS, and the resulting cell suspension was used to sort individual live cells in 96 well plates. Full length single cell RNA-seq was performed using a modified version of the Smart-seq2 protocol ^42^ as in ^65^. Briefly, individual cells were sorted into 96 well plates containing lysis buffer in presence of RNase inhibitor, dNTPs and OligodT. Reverse transcription of the polyadenylated RNA was performed with Maxima H Minus Reverse Transcriptase and Template Switching Oligos. The resulting cDNA was amplified with 23 cycles of PCR and libraries were prepared for sequencing with miniaturized NexteraXT Illumina protocol. Libraries were sequenced on Illumina NextSeq 500 System (single-end 75bp reads).

### Single cell RNA sequencing data processing

Following quality controls, sequencing reads were quality and adaptor trimmed using Trim Galore! v0.5.0 (https://www.bioinformatics.babraham.ac.uk/projects/trim_galore) (parameters: *--stringency 3 –q 20*). Trimmed reads were next aligned to the human reference genome (GRCh38.p13) using Hisat2^66^. Indexes for transcriptome-guided genome alignment were generated for the human GENCODE release 32 annotation (*extract_exons.py* and *extract_splice_sites.py* scripts). Gene expression levels were quantified with featureCounts v1.6.1 ^67^ (options: *-t exon -g gene_name*) using the human GENCODE release 32 annotation (https://www.gencodegenes.org/human/release_32.html). Multi-mapped reads were excluded from quantification. The following criteria were applied to exclude low-quality cells from subsequent analyses: less than 10,000 reads assigned to annotated transcripts; less than 2,000 detected genes, resulting in 394 cells.

### Single cell RNA sequencing data analysis

Single cell gene expression counts were next analyzed following the R/Bioconductor workflow ^68^. First, normalization factors were computed using the *computeSumFactors* function (*scran* package, default parameters) and read counts were normalized and log-transformed using the *logNormCounts* function (*scater* package, default parameters). Variance modeling for feature selection was next performed using the *modelGeneVar* function (*scran* package, default parameters). Each experimental batch was processed independently. The two experimental batches were then integrated using the matching Mutual Nearest Neighbors (MNN) approach ^69^ with the following pipeline: (i) selecting the top 4000 variable genes in each batch; (ii) running the *multiBatchNorm* function (*batchelor* package, default parameters) to harmonize library sizes across batches; running the *fastMNN* function (*batchelor* package, default parameters); using the top 50 MNN corrected dimensions for downstream analyses. Dimensionality reductions by t-SNE and UMAP were obtained using the *runTSNE* and *runUMAP* functions (*scater* package, parameters: perplexity = 10). Graph-based clustering was next performed using the *buildSNNGraph* function (*scran* package, parameters: k=10) and cluster_louvain function (*igraph* package). Cluster markers were identified by iteratively comparing gene expression levels of cells in each cluster against all the remaining cells (one vs all approach), implemented by running a Wilcoxon rank-sum test for each gene (*matrixTests* package), performing Bonferroni multiple hypothesis testing correction and reporting genes with adjusted p-value < 0.05 and expressed in at least 10% of cells in each comparison.

Pseudotime analysis was performed by re-running the batch correction pipeline described above for the selected cell types (settings: top 5000 variable genes and 50 top MNN components) and fitting a trajectory on the obtained UMAP space (*runUMAP* function from *scater* package, parameters: min_dist=0.05) with Monocle3 ^70^ (reverse graph embedding by running the *learn_graph* function with parameters and *order_cells* function with default parameters), setting as the starting point of the trajectory the earliest principal point of the 0: PSC 2D cells cluster.

For the comparative integrated analysis with the human embryo cell populations, the embryo dataset and the two experimental batches of the 3Depiblast dataset were integrated using the MNN approach with the following settings: (i) selecting the top 2500 variable genes in each batch; (ii) running the *multiBatchNorm*; running the *fastMNN* function. Clustering, cluster markers and t-SNE embedding (perplexity = 15) were then obtained as described above.

### Immunofluorescence analysis

3D structures were fixed with 2% Formaldehyde in PBS for 45 min at room temperature and then washed twice for 5 min in PBS. Permeabilisation and blocking were performed simultaneously by incubating samples with 1% BSA (Sigma Aldrich) in PBS with 0.5% Triton X-100 for 3 h at room temperature. Then samples were incubated 24 h at +4°C with primary antibodies (see Table 1), diluted in blocking solution. The day after, samples were washed with blocking solution three times for 15 min and then incubated 24 h at +4°C with secondary antibodies (see Table 1), diluted in blocking solution. Samples were then washed once with blocking solution for 15 min, then with PBS without MgCl2/CaCl2 two times for 15 min. Samples were adapted to and then mounted in Fluoromount-G (Southern Biotech). Nuclei were stained with Hoechst 33342 (ThermoFisher Cat. 62249) and F-actin with Phalloidin 488 (see Table 1). Images were acquired with a Leica SP5 confocal microscope equipped with a charge-coupled device camera usingLeica LAS AF software.

### Quantification of images

DAPI and phalloidin signals were quantified with FIJI software. We measure signal intensity along a line connecting the edge of each structure to the centre (radius). We took 7 measurements for each structure and analysed 3 structures for each sample, of similar size. Of each structure we analyse an equatorial confocal plane, containing a lumen, when present.

### Flow cytometry

After dissociation in single-cell suspension using TrypLE, cells were resuspended in PBS and filtered. Cells were analysed according to MIXL1-GFP expression with BD FACSCantoTM II cytometer and BD FACSDivaTM (v. 9.0) software.

### Quantitative PCR

Total RNA was isolated using Total RNA Purification Kit (Norgen Biotek 37500), and complementary DNA (cDNA) was made from 500 ng using M-MLV Reverse Transcriptase (Invitrogen 28025-013) and dN6 primers. For real-time PCR SYBR Green Master mix (Bioline BIO-94020) was used. Primers are detailed in Table 2. Three technical replicates were carried out for all quantitative PCR. GAPDH was used as an endogenous control to normalise expression. qPCR data were acquired with QuantStudio™ 6&7 Flex Software 1.0.

### Statistics and reproducibility

No statistical method was used to pre-determine sample size, but our sample sizes are similar to those reported in previous publications^13,47,49,51^. For each dataset, sample size *n* refers to experimental replicates and is represented by the number of dots in the plots or stated in the figure legends. Statistical analysis was performed within the R software environment (v4.0.4, https://www.R-project.org/). Statistical significance was determined using the two-tailed Welch’s unpaired *t*-test (*t.test* function with default parameters), or Quasi-Poisson regression for count data (*glm* function, family = “quasipoisson”). Data distribution was assumed to be normal, but this was not formally tested. No data points were excluded. P-values are reported in the plots, indicated as their numerical values. Unless stated otherwise, data are expressed as the mean ± standard errors of the mean (s.e.m.) of replicated independent experiments, shown as dots. All error bars indicate s.e.m.. All key experiments were repeated between two and four times independently, as indicated. All qPCR experiments were performed with three technical replicates.

### Data availability

The human embryo scRNA-seq dataset for comparative analyses was retrieved from the Gene Expression Omnibus (GEO) database, with accession code: GSE136447. The datasets generated in this study are available as raw data in the GEO database with accession code GSE248567.

#### ACKNOWLEDGMENTS

We thank all the members of the Martello Laboratory and Oliviero Laboratory for discussions and suggestions. In particular, we thank Martina Chieregato for technical support. We thank Stefania Rapelli, Isabelle Laurence Polignano and Fatemeh Mirzadeh for helping with the pilot experiment of single-cell RNA sequencing. We also thank Hannah T. Stuart for helping with immunostaining protocol in the 3D structures.

G.M.’s Laboratory is supported by grants from the Giovanni Armenise–Harvard Foundation, the Telethon Foundation (GJC21157), an ERC Starting Grant (MetEpiStem) and PRIN 2022. S.O. is supported by the Associazione Italiana per la Ricerca sul Cancro (AIRC) IG 2017 Id. 20240, and IG IG 2022 ID 27155, PRIN 2018, and IIGM institutional funds.

## AUTHOR CONTRIBUTIONS

I.Z. and E.C. set up the 3D-hE-gastruloid protocol, designed and performed most of the experiments.

I.Z. performed ZNF398 overexpression experiments. I.Z. and I.P. performed experiments with MIXL1-GFP reporter cells. I.Z. and A.D. performed molecular analyses. V.P. and I.Z. performed single-cell RNA sequencing experiments. A.L. performed analysis of single-cell RNA sequencing data and statistical analyses. I.Z., E.C., M.S. and A.L. prepared figures. M.A. analysed bulk RNA sequencing data. D.B. used the mouse ESC rosette assay. C.D. and G.A. designed and performed experiments with cysts composed of wild-type and Gata4-inducible ESCs. S.O. and G.M. supervised the study and secured fundings. G.M. performed image analysis and wrote the manuscript with suggestions from all authors.

## DECLARATION OF INTERESTS

The authors declare no competing interests.

---

**Extended Data Fig. 1.**
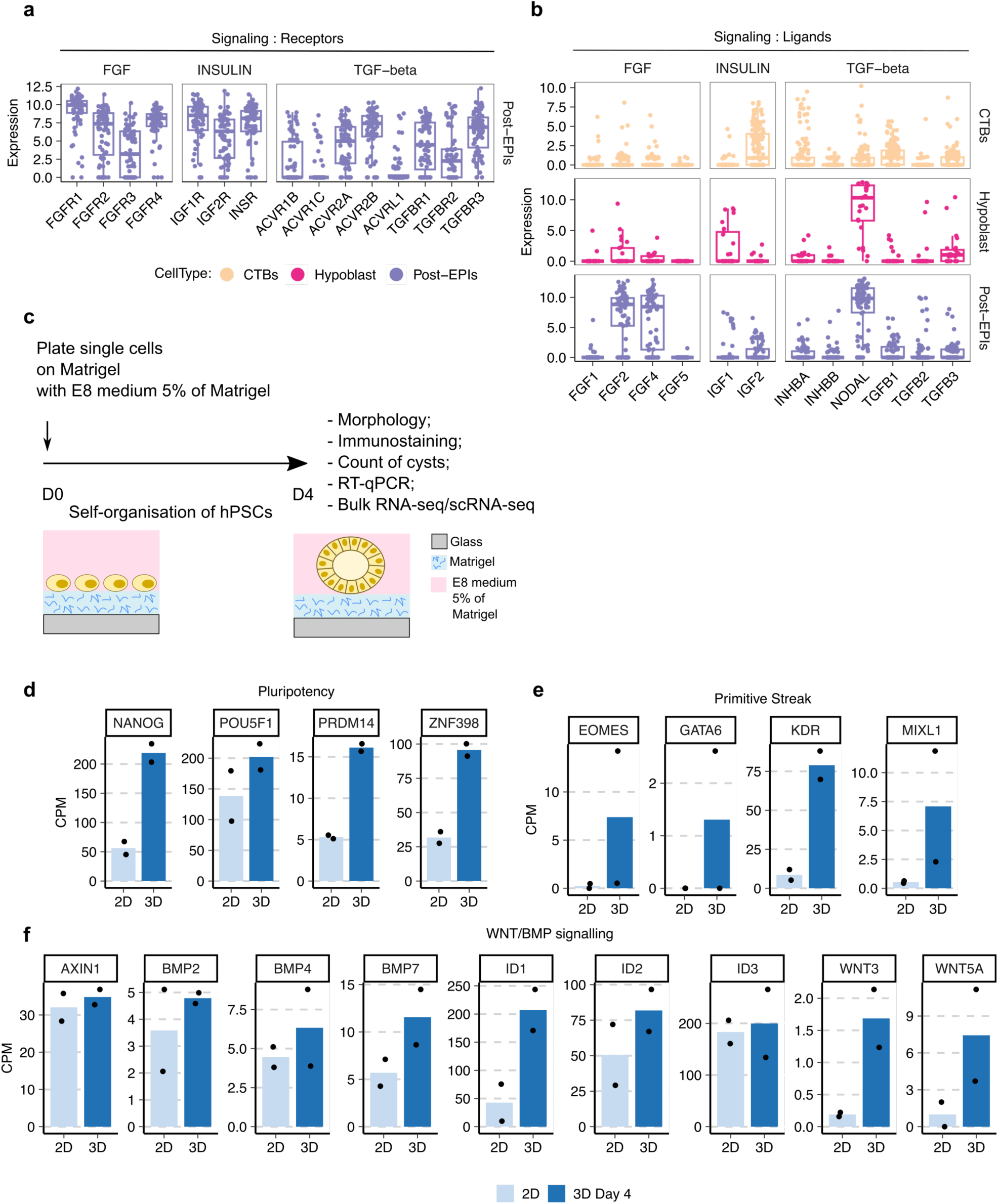
**a**, Box and dot plots showing the expression levels of receptors of the FGF, INSULIN and TGF-signalling pathways in the post-EPIs cell population of the embryo dataset^41^. Horizontal line indicates median, box indicates the interquartile range (IQR) and whiskers denote the 1.5 × IQR. **b,** Box and dot plots showing the expression levels of ligands of the FGF, INSULIN and TGF-signalling pathways in the post-EPIs, CTBs and Hypoblast cell population of the embryo dataset. Horizontal line indicates median, box indicates the interquartile range (IQR) and whiskers denote the 1.5 × IQR. **c,** Optimised protocol to mimic self-organisation *in vitro*. Single hPSCs were seeded on a Matrigel layer with E8 medium + 5% of Matrigel (v/v) After 4 days, 3D structures were analysed as indicated. See also Method details. **d-e-f,** Bar plots showing gene expression levels measured by RNAseq of KiPS cultured in the conventional 2D system or in 3D after 4 days of self-organisation for selected marker genes of pluripotency, primitive streak and Wnt/BMP signalling. Bars indicate the mean of 2 independent experiments, shown as dots.

**Extended Data Fig. 2.**
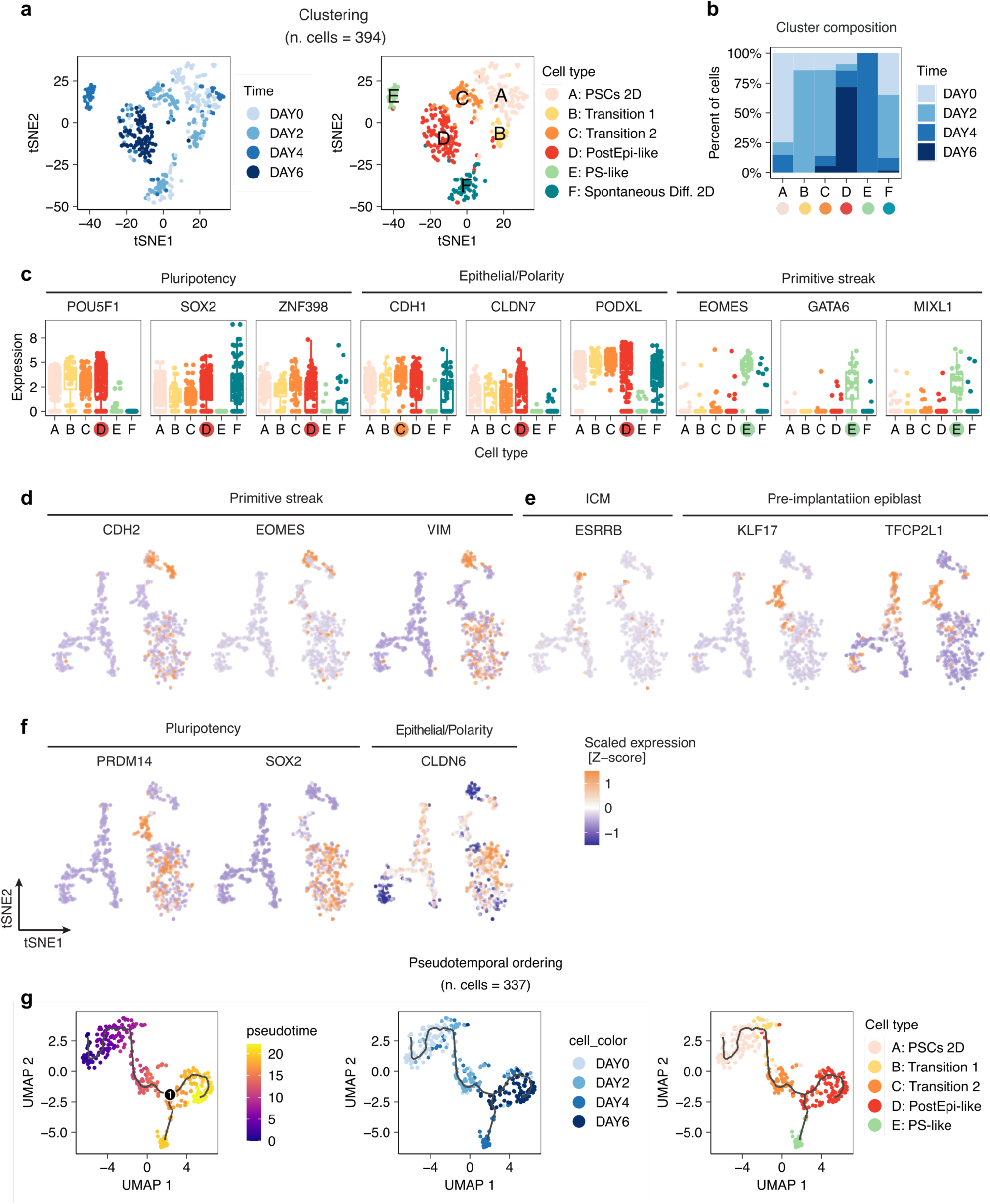
**a**, t-SNE embedding of 394 single cell transcriptomes showing the results of clustering analysis for the identification of cell populations in the 3D epiblast model dataset. Cells are coloured by their differentiation day (left) and inferred cell type (right). **b,** Barplots showing the relative proportions of cells by differentiation day in each cluster/inferred cell population. **c,** Box and dot plots showing the expression levels by cell type of selected marker genes for pluripotency, epithelial, polarity and primitive streak cell identities. Coloured dots indicate the cell cluster for which the gene was found as marker. **d-e-f,** t-SNE embedding of 892 single cell transcriptomes coloured according to the relative (Z-score) expression levels of selected marker genes for primitive streak (CDH2, EOMES, VIM) (E), ICM (ESRRB), Pre-implantation epiblast (KLF17, TFCP2L1) (F), pluripotency (PRDM14, SOX2) and epithelial/polarity (CLDN6) (G) cell identities. **g,** UMAP embedding of 337 single cell transcriptomes showing the results of pseudo-temporal ordering by reverse graph embedding using Monocle3. The line plot on the leftmost UMAP represents the embedded trajectory graph. Cells are coloured according to pseudotime (left), differentiation day (middle) and inferred cell type (right).

**Extended Data Fig. 3.**
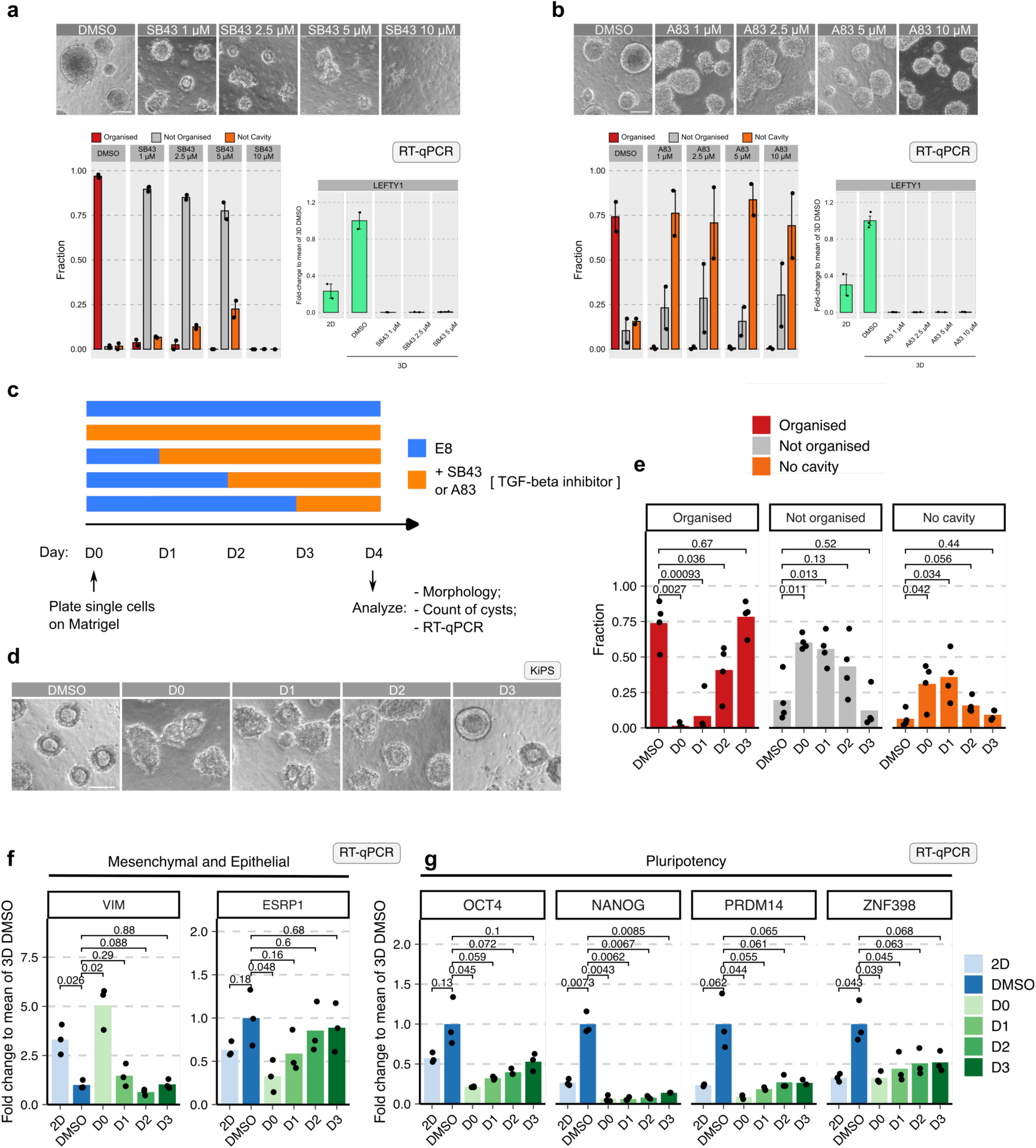
**a**, Top: Representative images of 3D structures treated with different doses of SB43. Bottom left: Bar plots showing the fraction of 3D structures in the indicated conditions. Bottom right: Bar plots showing gene expression analysis by RT-qPCR of LEFTY1 in the indicated conditions. **b,** Top: Representative images of 3D structures treated with different doses of A83. Bottom left: Bar plots showing the fraction of 3D structures in the indicated conditions. Bottom right: Bar plots showing gene expression analysis by RT-qPCR of LEFTY1 in the indicated conditions. **c,** Schematic representation of the experimental strategy used for TGF-beta inhibition. **d,** Representative images of KiPS-derived structures following a 4 days time-course of TGF-beta inhibition with SB43. **e,** Bar plots showing fraction of organised (red bars), not organised (grey bars) and no cavity (orange bars) KiPS-derived structures in a 4 days time-course of TGF-beta inhibition with SB43. Unpaired two-tailed Welch’s *t*-test. **f-g,** Bar plots showing gene expression analysis by RT-qPCR of mesenchymal/epithelial and polarity (left) and pluripotency (right) marker genes performed in KiPS-derived structures following a 4 days time-course of TGF-beta inhibition with SB43. Unpaired two-tailed Welch’s *t*-test.

**Extended Data Fig. 4.**
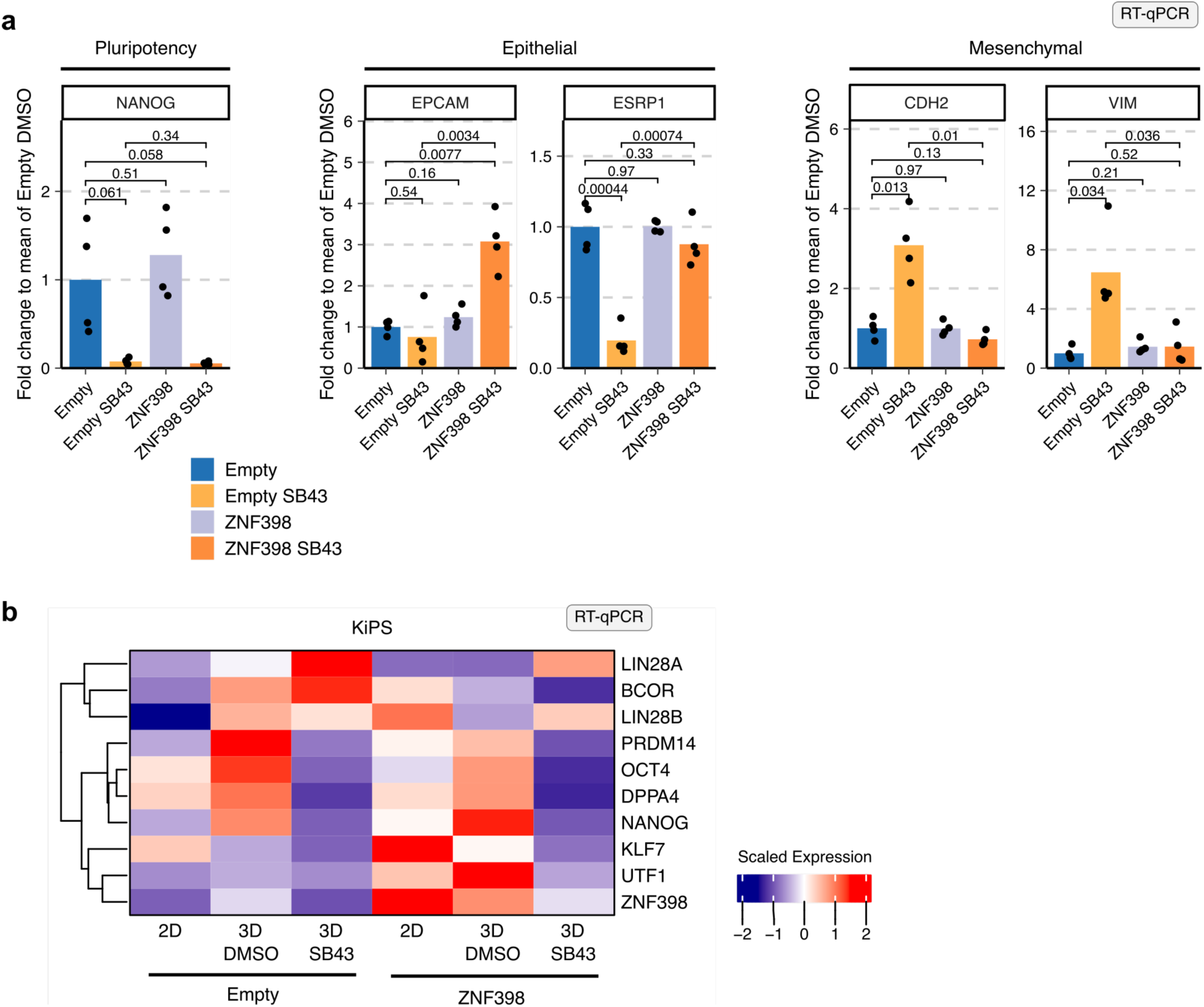
**a**, Gene expression analysis by qPCR of KiPS stably expressing the empty vector control (Empty) without (blue) or with SB43 (yellow), or expressing ZNF398 without SB43 (lilac) or with SB43 (orange) after 4 days of self-organisation. Bars indicate the mean of 4 independent experiments, shown as dots. Expression was normalised to the mean of Empty DMSO samples. Unpaired two-tailed Welch’s *t*-test. **b,** Heatmap showing gene expression analysis by qPCR for markers of pluripotency. Human PSCs, either expressing ZNF398 or an empty vector, were analysed when cultured under conventional conditions (2D) and after 4 days of self-organisation (3D) in presence of SB43 or DMSO. Values are expressed as fold-change relative to the mean of Empty 2D samples from 4 independent experiments and are scaled by rows (genes). Red and blue indicate high and low expression, respectively.

## Notes

### Competing Interest Statement

The authors have declared no competing interest.

## References

1. Boroviak, T. & Nichols, J. The birth of embryonic pluripotency. Philos Trans R Soc Lond B Biol Sci 369, 20130541 (2014).

2. Zhu, M. & Zernicka-Goetz, M. Principles of Self-Organization of the Mammalian Embryo. Cell 183, 1467–1478 (2020).

3. Pera, M. F. & Rossant, J. The exploration of pluripotency space: Charting cell state transitions in peri-implantation development. Cell Stem Cell 28, 1896–1906 (2021).

4. Rossant, J. & Tam, P. P. L. Early human embryonic development: Blastocyst formation to gastrulation. Developmental Cell 57, 152–165 (2022).

5. Sheng, G., Martinez Arias, A. & Sutherland, A. The primitive streak and cellular principles of building an amniote body through gastrulation. Science 374, abg1727 (2021).

6. Bedzhov, I. & Zernicka-Goetz, M. Self-Organizing Properties of Mouse Pluripotent Cells Initiate Morphogenesis upon Implantation. Cell 156, 1032–1044 (2014).

7. Kyprianou, C. et al. Basement membrane remodelling regulates mouse embryogenesis. Nature 582, 253–258 (2020).

8. Thomson, J. M., J. A. ItsKovitz-Eldor, J., Shapiro, S. S.,. Waknitz, M. A.,. Swierhiel, J. J.,. Marshall, V. S.,. Jones. Embryonic Stem Cell Lines Derived from Human Blastocysts. Science (1998) doi:10.1126/science.282.5391.1145.

9. Takahashi, K. et al. Induction of Pluripotent Stem Cells from Adult Human Fibroblasts by Defined Factors. Cell 131, 861–872 (2007).

10. Yu, J. et al. Induced pluripotent stem cell lines derived from human somatic cells. Science 318, 1917–1920 (2007).

11. Tyser, R. C. V. et al. Single-cell transcriptomic characterization of a gastrulating human embryo. Nature 600, 285–289 (2021).

12. Wang, Z., Oron, E., Nelson, B., Razis, S. & Ivanova, N. Distinct lineage specification roles for NANOG, OCT4, and SOX2 in human embryonic stem cells. Cell Stem Cell 10, 440–454 (2012).

13. Zorzan, I. et al. The transcriptional regulator ZNF398 mediates pluripotency and epithelial character downstream of TGF-beta in human PSCs. Nature Communications 11, 2364 (2020).

14. Chia, N. Y. et al. A genome-wide RNAi screen reveals determinants of human embryonic stem cell identity. Nature 468, 316–320 (2010).

15. Eiselleova, L. et al. A complex role for FGF-2 in self-renewal, survival, and adhesion of human embryonic stem cells. Stem Cells 27, 1847–1857 (2009).

16. Greber, B. et al. Conserved and divergent roles of FGF signaling in mouse epiblast stem cells and human embryonic stem cells. Cell Stem Cell 6, 215–226.

17. Martello, G. & Smith, A. The Nature of Embryonic Stem Cells. Annual Review of Cell and Developmental Biology 30, 647–675 (2014).

18. Singh, A. M. et al. Signaling Network Crosstalk in Human Pluripotent Cells: A Smad2/3-Regulated Switch that Controls the Balance between Self-Renewal and Differentiation. Cell Stem Cell 10, 312–326 (2012).

19. Vallier, L. Activin/Nodal and FGF pathways cooperate to maintain pluripotency of human embryonic stem cells. Journal of Cell Science 118, 4495–4509 (2005).

20. Vallier, L. et al. Activin/Nodal signalling maintains pluripotency by controlling Nanog expression. Development 136, 1339–1349 (2009).

21. Wamaitha, S. E. et al. IGF1-mediated human embryonic stem cell self-renewal recapitulates the embryonic niche. Nature Communications 11, 1–16 (2020).

22. Ai, Z. et al. Dissecting peri-implantation development using cultured human embryos and embryo-like assembloids. 2023.06.15.545180 Preprint at 10.1101/2023.06.15.545180 (2023).

23. Hislop, J. et al. Modelling Human Post-Implantation Development via Extra-Embryonic Niche Engineering. 2023.06.15.545118 Preprint at 10.1101/2023.06.15.545118 (2023).

24. Liu, L. et al. Modeling post-implantation stages of human development into early organogenesis with stem-cell-derived peri-gastruloids. Cell 0, (2023).

25. Moris, N. et al. An in vitro model of early anteroposterior organization during human development. Nature 582, 410–415 (2020).

26. Oldak, B. et al. Transgene-Free Ex Utero Derivation of A Human Post-Implantation Embryo Model Solely from Genetically Unmodified Naïve PSCs. 2023.06.14.544922 Preprint at 10.1101/2023.06.14.544922 (2023).

27. Pedroza, M. et al. Self-patterning of human stem cells into post-implantation lineages. Nature 1–3 (2023) doi:10.1038/s41586-023-06354-4.

28. Simunovic, M., Siggia, E. D. & Brivanlou, A. H. In vitro attachment and symmetry breaking of a human embryo model assembled from primed embryonic stem cells. Cell Stem Cell 29, 962–972.e4 (2022).

29. Simunovic, M. et al. A 3D model of a human epiblast reveals BMP4-driven symmetry breaking. Nature Cell Biology 21, 900 (2019).

30. Sozen, B. et al. Reconstructing aspects of human embryogenesis with pluripotent stem cells. Nat Commun 12, 5550 (2021).

31. Weatherbee, B. A. T. et al. A model of the post-implantation human embryo derived from pluripotent stem cells. Nature 1–3 (2023) doi:10.1038/s41586-023-06368-y.

32. Zheng, Y. et al. Controlled modelling of human epiblast and amnion development using stem cells. Nature 1–5 (2019) doi:10.1038/s41586-019-1535-2.

33. Taniguchi, K. et al. Lumen Formation Is an Intrinsic Property of Isolated Human Pluripotent Stem Cells. Stem Cell Reports 5, 954–962 (2015).

34. Shao, Y. et al. Self-organized amniogenesis by human pluripotent stem cells in a biomimetic implantation-like niche. Nature Materials 16, 419–427 (2017).

35. Baillie-Benson, P., Moris, N. & Martinez Arias, A. Pluripotent stem cell models of early mammalian development. Current Opinion in Cell Biology 66, 89–96 (2020).

36. Shahbazi, M. N. et al. Self-organization of the human embryo in the absence of maternal tissues. Nat Cell Biol 18, 700–708 (2016).

37. Shao, Y. et al. Self-organized amniogenesis by human pluripotent stem cells in a biomimetic implantation-like niche. Nature Materials 16, 419–427 (2017).

38. Ludwig, T. E. et al. Derivation of human embryonic stem cells in defined conditions. Nature Biotechnology 24, 185–187 (2006).

39. Chen, G. et al. Chemically defined conditions for human iPS cell derivation and culture. Nat Methods 8, 424–429 (2011).

40. Deglincerti, A. et al. Self-organization of the in vitro attached human embryo. Nature 533, 251–254 (2016).

41. Xiang, L. et al. A developmental landscape of 3D-cultured human pre-gastrulation embryos. Nature 577, 537–542 (2020).

42. Picelli, S. et al. Full-length RNA-seq from single cells using Smart-seq2. Nature Protocols 9, 171–181 (2014).

43. Molè, M. A. et al. A single cell characterisation of human embryogenesis identifies pluripotency transitions and putative anterior hypoblast centre. Nat Commun 12, 3679 (2021).

44. Stirparo, G. G. et al. Integrated analysis of single-cell embryo data yields a unified transcriptome signature for the human pre-implantation epiblast. Development 145, dev158501 (2018).

45. Matthews, K. R. W., Wagner, D. S. & Warmflash, A. Stem Cell-Based Models of Embryos: The Need for Improved Naming Conventions. Stem Cell Reports (2021) doi:10.1016/j.stemcr.2021.02.018.

46. Yoney, A. et al. WNT signaling memory is required for ACTIVIN to function as a morphogen in human gastruloids. eLife 7, e38279 (2018).

47. Shahbazi, M. N. et al. Pluripotent state transitions coordinate morphogenesis in mouse and human embryos. Nature 552, 239–243 (2017).

48. Bedzhov, I. & Zernicka-Goetz, M. Self-Organizing Properties of Mouse Pluripotent Cells Initiate Morphogenesis upon Implantation. Cell 156, 1032–1044 (2014).

49. Carbognin, E. et al. Esrrb guides naive pluripotent cells through the formative transcriptional programme. Nature Cell Biology 25, 643–657 (2023).

50. Papanayotou, C. et al. A Novel Nodal Enhancer Dependent on Pluripotency Factors and Smad2/3 Signaling Conditions a Regulatory Switch During Epiblast Maturation. PLOS Biology 12, e1001890 (2014).

51. Amadei, G. et al. Inducible Stem-Cell-Derived Embryos Capture Mouse Morphogenetic Events In Vitro. Developmental Cell 56, 366–382.e9 (2021).

52. Bernardo, A. S. et al. BRACHYURY and CDX2 mediate BMP-induced differentiation of human and mouse pluripotent stem cells into embryonic and extraembryonic lineages. Cell Stem Cell 9, 144–155 (2011).

53. McLean, A. B. et al. Activin a efficiently specifies definitive endoderm from human embryonic stem cells only when phosphatidylinositol 3-kinase signaling is suppressed. Stem Cells 25, 29–38 (2007).

54. Sturgeon, C. M., Ditadi, A., Awong, G., Kennedy, M. & Keller, G. Wnt Signaling Controls the Specification of Definitive and Primitive Hematopoiesis From Human Pluripotent Stem Cells. Nat Biotechnol 32, 554–561 (2014).

55. D’Amour, K. A. et al. Efficient differentiation of human embryonic stem cells to definitive endoderm. Nature Biotechnology 23, 1534–1541 (2005).

56. Funa, N. S. et al. β-Catenin Regulates Primitive Streak Induction through Collaborative Interactions with SMAD2/SMAD3 and OCT4. Cell Stem Cell 16, 639–652 (2015).

57. Amadei, G. et al. Embryo model completes gastrulation to neurulation and organogenesis. Nature 610, 143–153 (2022).

58. Heldin, C.-H., Landström, M. & Moustakas, A. Mechanism of TGF-beta signaling to growth arrest, apoptosis, and epithelial-mesenchymal transition. Curr Opin Cell Biol 21, 166–176 (2009).

59. Hikasa, H. & Sokol, S. Y. Wnt Signaling in Vertebrate Axis Specification. Cold Spring Harb Perspect Biol 5, a007955 (2013).

60. Warmflash, A., Sorre, B., Etoc, F., Siggia, E. D. & Brivanlou, A. H. A method to recapitulate early embryonic spatial patterning in human embryonic stem cells. Nature Methods 11, 847–854 (2014).

61. Beyer, T. A. et al. Switch enhancers interpret TGF-β and hippo signaling to control cell fate in human embryonic stem cells. Cell Reports 5, 1611–1624 (2013).

62. Takashima, Y. et al. Resetting Transcription Factor Control Circuitry toward Ground-State Pluripotency in Human. Cell 158, 1254–1269 (2014).

63. Davis, R. P. et al. Targeting a GFP reporter gene to the MIXL1 locus of human embryonic stem cells identifies human primitive streak–like cells and enables isolation of primitive hematopoietic precursors. Blood 111, 1876–1884 (2008).

64. Love, M. I., Huber, W. & Anders, S. Moderated estimation of fold change and dispersion for RNA-seq data with DESeq2. Genome Biology 15, 550 (2014).

65. Proserpio, V., Duval, C., Falvo, V., Donati, G. & Oliviero, S. Single-Cell Sequencing for Everybody. in Immune Receptors: Methods and Protocols (eds. Rast, J. & Buckley, K.) 217–229 (Springer US, 2022). doi:10.1007/978-1-0716-1944-5_15.

66. Kim, D., Paggi, J. M., Park, C., Bennett, C. & Salzberg, S. L. Graph-based genome alignment and genotyping with HISAT2 and HISAT-genotype. Nat Biotechnol 37, 907–915 (2019).

67. Liao, Y., Smyth, G. K. & Shi, W. featureCounts: an efficient general purpose program for assigning sequence reads to genomic features. Bioinformatics 30, 923–930 (2014).

68. Amezquita, R. A. et al. Orchestrating single-cell analysis with Bioconductor. Nat Methods 17, 137–145 (2020).

69. Haghverdi, L., Lun, A. T. L., Morgan, M. D. & Marioni, J. C. Batch effects in single-cell RNA-sequencing data are corrected by matching mutual nearest neighbors. Nat Biotechnol 36, 421–427 (2018).

70. Cao, J. et al. The single-cell transcriptional landscape of mammalian organogenesis. Nature 566, 496–502 (2019).

