## Supplementary Tables 1 and 2 for "TGF-beta dynamically controls epithelial identity in a 3D model of human epiblast"

Table 1. Antibodies details

Table 2. qPCR primers details

**Table 1:** Antibodies details

| **Antibody** | **Dilution** | **Product code** | **Company** |
| --- | --- | --- | --- |
| OCT4 | 1:300 | sc-5279 | Santa Cruz Biotechnologies |
| PODXL | 1:200 | MAB1556 | R&D |
| SOX2 | 1:500 | NB11037235 | Novus |
| T | 1:100 | AF2085 | R&D |
| Alexa Fluor 488 Phalloidin | 1:200 | A12379 | ThermoFisher |
| Donkey anti-Goat IgG (H+L) Secondary Antibody Alexa Fluor 488 | 1:500 | A11055 | ThermoFisher |
| Donkey anti-Rabbit IgG (H+L) Secondary Antibody Alexa Fluor 568 | 1:500 | A10042 | ThermoFisher |

**Table 2**: qPCR primers details

| Gene | Forward | Reverse |
| --- | --- | --- |
| BCOR | CTCAGGGGCCACGAAACT | ATGGTGACAATTTCCACGTTT |
| DPPA4 | TCTGGTGTCAGGTGGTGTGT | TCCCTTCTTGCTTTTCTGGA |
| EOMES | CAACATAAACGGACTCAATCCCA | ACCACCTCTACGAACACATTGT |
| EPCAM | TGGACATAGCTGATGTGGCTTA | CCAGGATCCAGATCCAGTTG |
| ESRP1 | CTTCCAACCCCTCCCATTAT | ATTGTGGCTGCATAGGGAAG |
| FLK1 | GGCGGCACGAAATATCCTCT | GGAGGCGAGCATCTCCTTTT |
| GAPDH | CGAGATCCCTCCAAAATCAA | GGCAGAGATGATGACCCTTT |
| GATA4 | GGAAGCCCAAGAACCTGAAT | GTTGCTGGAGTTGCTGGAA |
| GATA6 | GCAAAAATACTTCCCCCACA | TCTCCCGCACCAGTCATC |
| KLF7 | CTCATGGGAGGGATGTGAGT | ACCTGGAAAAACACCTGTCG |
| LEFTY1 | GGACCTTGGGGACTATGGAG | ATCCCCTGCAGGTCAATGTA |
| LIN28A | CTGTAAGTGGTTCAACGTGCG | CCATGTGCAGCTTACTCTGGT |
| LIN28B | CCTCCTCAGCCAAAGAAGTG | TGGGATTCTGCTTCCTGTCT |
| MIXL1 | GGTACCCCGACATCCACTTG | TAATCTCCGGCCTAGCCAAA |
| NANOG | TTTGTGGGCCTGAAGAAAACT | AGGGCTGTCCTGAATAAGCAG |
| NCAD/CDH2 | CGTGGTCAAACCAATCGAC | AACAGACACGGTTGCAGTTG |
| OCT4 | GTGGAGGAAGCTGACAACAA | ATTCTCCAGGTTGCCTCTCA |
| PRDM14 | GAGCCTTCAGGTCACAGAGC | TCCACACAGGGGGTGTACTT |
| SOX2 | CCAGCAGACTTCACATGTCC | ACATGTGTGAGAGGGGCAGT |
| SOX17 | ACGCCGAGTTGAGCAAGA | TCTGCCTCCTCCACGAAG |
| T | TATGAGCCTCGAATCCACATAGT | CCTCGTTCTGATAAGCAGTCAC |
| UTF1 | CTCCCAGCGAACCAGACG | GGAGGCGTCCGCAGACTT |
| VIM | CTCCACGAAGAGGAAATCCA | GTGAGGTCAGGCTTGGAAAC |
| ZNF398 | TGGCAAGAATCTCAGCCAAGA | GTGGAGTAAAGTGCTTAGGGC |
| Gapdh | ATCCTGCACCACCAACTGCT | GGGCCATCCACAGTCTTCTG |
| Oct4 | GTTGGAGAAGGTGGAACCAA | CTCCTTCTGCAGGGCTTTC |
